# CTP and *parS* coordinate ParB partition complex dynamics and ParA-ATPase activation for ParABS-mediated DNA partitioning

**DOI:** 10.1101/2021.01.24.427996

**Authors:** James A. Taylor, Yeonee Seol, Jagat Budhathoki, Keir C. Neuman, Kiyoshi Mizuuchi

## Abstract

ParABS partition systems, comprising the centromere-like DNA sequence *parS,* the *parS*-binding ParB-CTPase and the nucleoid-binding ParA-ATPase, ensure faithful segregation of bacterial chromosomes and low-copy-number plasmids. F-plasmid partition complexes containing ParB_F_ and *parS_F_* move by generating and following a local concentration gradient of nucleoid-bound ParA_F_. However, the process through which ParB_F_ activates ParA_F_-ATPase has not been defined. We studied CTP- and *parS_F_*-modulated ParA_F_—ParB_F_ complex assembly, in which DNA-bound ParA_F_-ATP dimers are activated for ATP hydrolysis by interacting with two ParB_F_ N-terminal domains. CTP or *parS_F_* enhances the ATPase rate without significantly accelerating ParA_F_—ParB_F_ complex assembly. Together, *parS_F_* and CTP accelerate ParA_F_—ParB_F_ assembly without further significant increase in ATPase rate. Magnetic-tweezers experiments showed that CTP promotes multiple ParB_F_ loading onto *parS_F_*-containing DNA, generating condensed partition complex-like assemblies. We propose that ParB_F_ in the partition complex adopts a conformation that enhances ParB_F_—ParB_F_ and ParA_F_—ParB_F_ interactions promoting efficient partitioning.

## Introduction

Faithful segregation of replicated chromosomes is essential for efficient proliferation of cells. Accordingly, many bacteria are equipped with active chromosome and plasmid partition systems belonging to the ParABS family (Baxter and Funnell, 2014; Lutkenhaus, 2012; Vecchiarelli et al., 2012). Basic ParABS systems comprise two proteins, ParA and ParB, and a centromere-like, cis-acting DNA element called *parS*. The ParA proteins of this family are ATPases with a characteristic deviant Walker-A motif (Motallebi-Veshareh et al., 1990) and bind non-specific DNA (nsDNA) in an ATP-dependent manner by forming a DNA binding-competent dimer (Davey and Funnell, 1994; Leonard et al., 2005; Vecchiarelli et al., 2010). Accordingly, ParA proteins localize to the bacterial chromosome (the nucleoid) *in vivo* (Ebersbach and Gerdes, 2004; Hatano et al., 2007; Lim et al., 2014).

ParB is typically a dimeric sequence-specific DNA binding protein that binds tightly to the *parS* consensus sequences that mark the DNA cargo to be partitioned (Bouet et al., 2000; Mori et al., 1989; Pillet et al., 2011; Taylor et al., 2015). Most ParABS systems have multiple copies of a ParB dimer binding consensus sequence that collectively constitute a *parS* site. F-plasmid has a *parS* sequence cluster (*parS_F_*, also called *sopC*) composed of twelve repeats of a 16 bp consensus sequence, each separated by 27 base-pair spacer sequences (Helsberg and Eichenlaub, 1986). The ParBs of known chromosomal and plasmid Par systems such as P1 and F bind to *parS via* a helix-turn-helix motif (Schumacher and Funnell, 2005; Schumacher et al., 2010). These HTH-ParB proteins also associate with several kilobases of DNA surrounding a *parS* site *in vivo* in a proximity-dependent manner without obvious sequence specificity (Breier and Grossman, 2007; Murray et al., 2006; Rodionov et al., 1999; Sanchez et al., 2015). This activity, known as ParB spreading, is believed to be essential for proper function of these systems (Breier and Grossman, 2007; Graham et al., 2014) and results in the formation of a large nucleo-protein complex (the partition complex) around the *parS* site on the DNA to be partitioned. Mutations in the *B. subtilis parB* gene blocking spreading and causing partition deficiency have been identified within the Box II region (GXRR) of the N-terminal domain (Breier and Grossman, 2007; Graham et al., 2014), a highly conserved motif among HTH-ParB homologues (Yamaichi and Niki, 2000). Recently, several groups reported that HTH-ParB proteins have CTPase activity and the Box II residues play key roles in CTP binding and hydrolysis, suggesting that ParB spreading is driven by an active process dependent on energy derived from CTP hydrolysis (Jalal et al., 2020; Osorio-Valeriano et al., 2019; Soh et al., 2019).

ParB interacts with ParA *via* its N-terminal region (Ravin et al., 2003) and activates nsDNA-bound ParA dimer’s ATPase, releasing it from DNA (Ah-Seng et al., 2009; Davis et al., 1992; Scholefield et al., 2011; Watanabe et al., 1992). In the absence of ParB stimulation, the ATP turnover of ParA is low, typically around one ATP per hour (Ah-Seng et al., 2009; Davis et al., 1992; Fung et al., 2001; Scholefield et al., 2011). Because of this slow basal ATPase rate, and since the majority of cellular ParB molecules is concentrated at the partition complexes due to ParB spreading, ATP hydrolysis by ParA and dissociation from the nucleoid is expected to occur principally in the vicinity of partition complexes. Biochemical studies of P1 ParA ATPase showed a significant time-delay before activation of ParA for DNA binding after ATP binding, predicting a significant free bulk-diffusion period for ParA before reactivation for nsDNA binding (Vecchiarelli et al., 2010). This, along with *in vivo* imaging observations (Hatano et al., 2007; Ringgaard et al., 2009), led to a prediction that the nucleoid proximal to a partition complex would become depleted of ParA (the ParA depletion zone) and the proposal of a diffusion-ratchet model for plasmid segregation by the ParABS system (Vecchiarelli et al., 2010).

The diffusion-ratchet model is based on the premise that the nucleoid-bound ParA-ATPase activation by plasmid-bound ParB generates a local ParA depletion zone on the nucleoid, forming a nucleoid-bound ParA concentration gradient around the partition complex (Hu et al., 2017; Vecchiarelli et al., 2010). The interaction of plasmid-bound ParB with the ParA gradient on the nucleoid results in a cargo position-dependent free-energy difference (Sugawara and Kaneko, 2011). Binding of ParB to ParA reduces the system free-energy, therefore moving the cargo to a higher ParA concentration lowers the system free-energy. This cargo position-dependent free-energy difference translates to a directional motive force on the ParB bound cargo. The generation of sufficient cargo motive force to overcome thermal diffusion was demonstrated in cell-free reconstitution experiments showing that a bead coated with *parS_F_*-containing DNA is driven across an nsDNA-coated flow cell surface in the presence of ParA_F_, ParB_F_ and ATP (Vecchiarelli et al., 2014). In some *in vivo* time-lapse imaging experiments ParA has been observed to undergo pole-to-pole oscillations along the length of a nucleoid with a partition complex chasing the receding edge of a ParA distribution zone on the nucleoid, further supporting the diffusion-ratchet model (Hatano et al., 2007; Ringgaard et al., 2009). Variations of this model have also been proposed (Le Gall et al., 2016; Lim et al., 2014; McLeod et al., 2017).

Generating persistent directional motion by the diffusion-ratchet model requires a balance between the ParA—ParB association/dissociation dynamics prior to ATP hydrolysis and the steps and kinetic parameters that govern ATP hydrolysis. For example, if every ParA—ParB association resulted in instantaneous ATP hydrolysis and dissolution of the ParA—ParB bonds linking the partition complex (cargo) and the nucleoid, no cargo driving force would result. Conversely, if each bond persisted too long the cargo would remain immobile on the nucleoid (Hu et al., 2017). However, the detailed biochemical reaction steps leading to ParB activation of DNA-bound ParA-ATPase and the subsequent release of ParA have not been determined.

In order to advance our quantitative understanding of the ParABS system mechanism, we investigated the assembly and disassembly kinetics of ParA_F_—ParB_F_ complexes that form prior to ATP hydrolysis using F-plasmid ParA_F_B_F_S_F_ (also called SopA/B/C) as a model system. Employing a TIRF microscopy-based nsDNA-carpet assay (Vecchiarelli et al., 2013) we first examined the stoichiometry of the nsDNA-bound ParA_F_—ParB_F_ complexes that accumulate in the absence of ATP hydrolysis. We investigated impacts of different ParB_F_-cofactors (*parS_F_* and CTP) or ParB_F_ mutations that hinder ParB_F_ dimerization, *parS_F_*-binding, CTPase activity, or ParA_F_ ATPase activation. We then studied how the same set of cofactors or ParB_F_ mutations influenced ParA_F_-ATPase activation by ParB_F_. Our results showed that ParB_F_ formed complexes with nsDNA-bound ParA_F_ in which both ParB_F_-interacting faces of the ParA_F_ dimers were occupied by the N-terminal ParA_F_-activation domain of ParB_F_. All such complexes observed in the presence of ATP*γ*S exhibited similar, slow dissociation kinetics from nsDNA compared to ParA_F_ dimers in the absence of ParB_F_ so long as both ParB_F_-interacting faces of the ParA_F_ dimers were occupied by ParB_F_ N-terminal domains. Binding of ParB_F_ N-terminal domains at both ParB_F_-interaction faces is also necessary for efficient ATPase activation of nsDNA-bound ParA_F_ dimers. Strikingly, the ParB_F_ cofactors, CTP and *parS_F_*, acted synergistically to accelerate the assembly of pre-ATP hydrolysis ParB_F_—ParA_F_ complexes on nsDNA. In addition, a magnetic tweezers-based DNA condensation assay revealed that stable DNA condensation by ParB_F_ required both CTP and *parS_F_.* These observations suggested that CTP and *parS_F_* promote both ParA_F_—ParB_F_ and ParB_F_—ParB_F_ interactions. The compaction of *parS_F_*-containing DNA by ParB_F_ in the presence of CTP recapitulated the salient features of the condensed ParB spreading partition complex observed *in vivo*, suggesting that ParB_F_ in the presence of *parS_F_* and CTP closely reflects the functional state of ParB_F_ in the partition complexes *in vivo*. Interestingly, ParB_F_ in these conditions accelerated ATP turnover by the nsDNA-bound ParA_F_ no more than two-fold compared to ParB_F_ without *parS_F_* and CTP, to a modest ∼80 h^-1^. These findings have important implications for the diffusion-ratchet model of F-plasmid partition by the ParAB*S* system.

## Results

Here, we investigated the ParA—ParB interactions that lead to accelerated ATP hydrolysis by ParA and how they are impacted by *parS_F_* and CTP. First, we addressed the nature of nsDNA-bound ParA_F_—ParB_F_ complexes. In the presence of ATP, ParA_F_ forms DNA binding-competent dimers and binds the nsDNA-carpet without ParB_F_. Upon forming a complex with ParB_F_, ParA_F_ becomes activated for ATP hydrolysis and the complex rapidly disassembles (Vecchiarelli et al., 2013), impeding characterization of the complex. Therefore, we studied DNA-bound ParA_F_— ParB_F_ complexes that accumulate prior to ATP hydrolysis by using non-hydrolysable ATPγS. ParA_F_ is not expected to form DNA binding competent dimers efficiently in the presence of ATPγS based on the study of the closely related ParA_P1_ ATPase (Vecchiarelli et al., 2010). However, we found ParB_F_ promotes conversion of ParA_F_ to a DNA binding competent state in the presence of ATPγS (see below). Hence in this study, we infused ParA_F_-eGFP and ParB_F_-Alexa647, or other fluorescent-labeled ParB_F_ variants, preincubated at room temperature for 10 min with ATPγS into an nsDNA-carpeted flow cell and quantified the densities of the two DNA-associated proteins by imaging the flow cell surface with TIRF microscopy (Figure 1). ParA_F_ alone at the concentration used here (1 μM, all protein concentrations are expressed as monomer concentrations) does not efficiently bind DNA in the presence of ATPγS as expected, and only low-level steady state density (less than ∼ 200 monomers per μm^2^) of ParA_F_-eGFP was detected on the DNA-carpet (Figure 2A, B). The observed DNA dissociation rate constant of ParA_F_-ATPγS (∼6 min^-1^) is similar to that of ParA_F_-ATP estimated by FRAP, or by washing the flow-cell with nsDNA-containing buffer (∼4.5 – 6 min^-1^, Vecchiarelli, 2013). ParB_F_-Alexa647 (2 μM) did not bind the DNA-carpet to a significant level in the 300 mM K-glutamate buffer used in this experiment.

**Figure 1.**
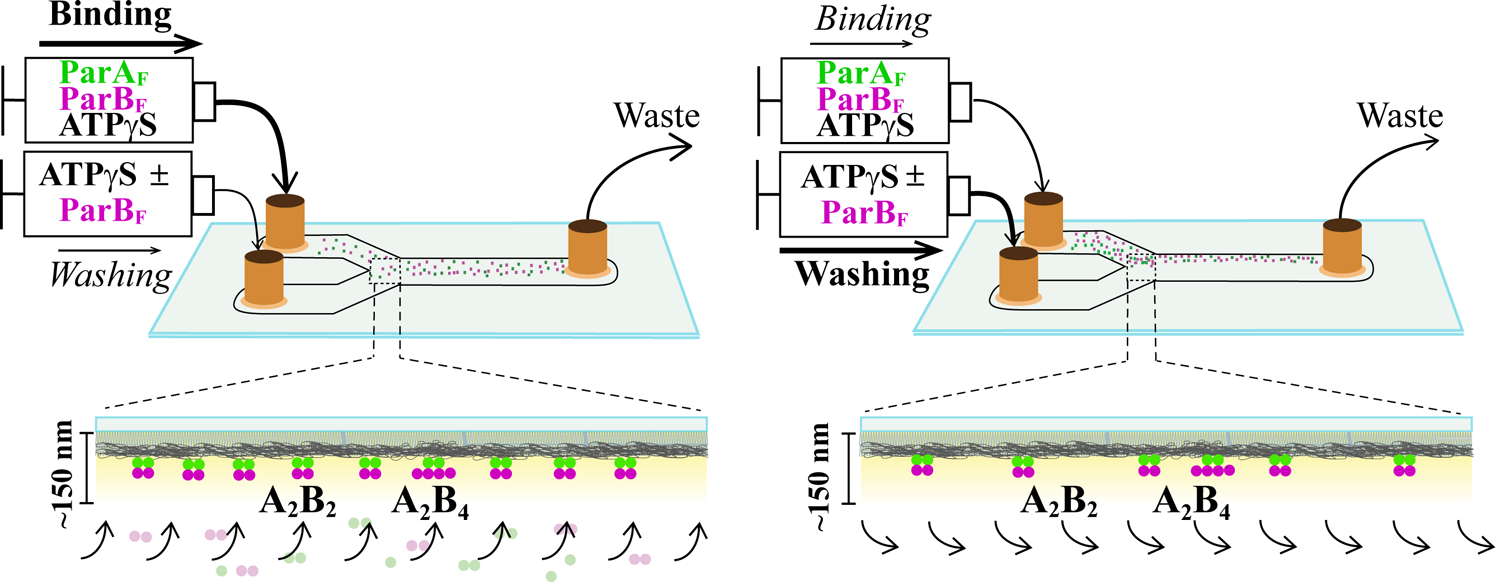
Schematic of flow cell setup for visualizing the binding and dissociation of fluorescent proteins on DNA-carpet. ParA_F_-eGFP and ParB_F_-Alexa647 proteins were flowed over a dense carpet of nsDNA attached to the supported lipid bilayer coated surface of a flow cell. TIRF microscopy permits selective detection of the DNA-carpet bound proteins. Sample solution and wash buffer, as specified for each experiment, were infused *via* two syringes at different infusion rates from separate inlets into a Y-shaped flow cell. A laminar boundary separates the two solutions downstream of the flow convergence point at the Y-junction. At the midpoint across the flow channel, downstream but close to the flow convergence point where the observations are made, the DNA-carpet area is exposed to the syringe content of the higher infusion rate. When the infusion rates of the two syringes are switched, the laminar boundary moves across the observation area and the solution flowing over the area switches. By switching the infusion rates of the two syringes repeatedly, multiple DNA-carpet-bound protein complex assembly and wash cycles can be recorded.

**Figure 2.**
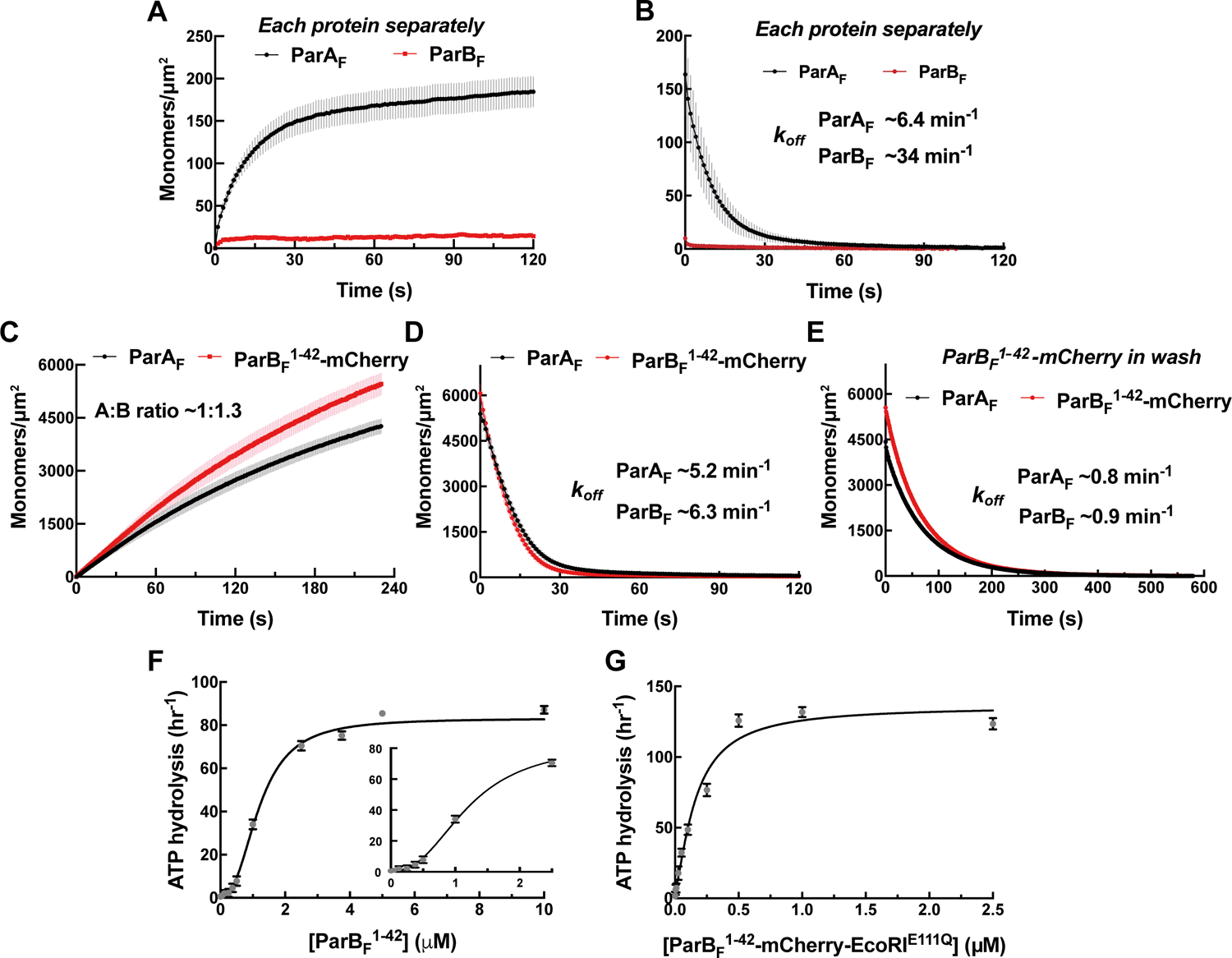
Monomeric ParB_F_^1-42^-mCherry can activate ParA_F_ for nsDNA binding in the presence of ATPγS by forming a ∼1:1 complex. Protein sample solution in the presence of ATPγS (1 mM) was infused into nsDNA-carpeted flow cell at a constant flow to monitor the protein binding to the nsDNA, and the sample solution was switched to a wash buffer containing ATPγS to monitor protein dissociation from nsDNA. (A and B) Binding to, and dissociation from, nsDNA of ParA_F_-eGFP (1 μM) or ParB_F_-Alexa647 (2 μM) were measured separately. (C) ParB_F_^1-42^-mCherry (10 μM) and ParA_F_-eGFP (1 μM) preincubated with ATPγS were infused into the nsDNA-carpeted flow cell and (D) washed with buffer containing ATPγS. (E) The washing experiment of D was repeated with wash buffer containing ATPγS and ParB_F_^1-42^-mCherry (10 μM). For the parameters of the time courses of above experiments and subsequent experiments of the same type in this study, see Table 1. The ParB_F_:ParA_F_ ratio was calculated from carpet-bound densities of the two proteins measured in parallel, and summarized in Table 1. (F) ParA_F_-ATPase activity (expressed as turnover rate per ParA_F_ monomer) was measured in the presence of EcoRI-digested pBR322 DNA (60 μg/ml) and different concentrations of ParB_F_^1-42^. Inset shows a plot with expanded abscissa. (G) ParA_F_-ATPase activity was measured as above in the presence of dimeric ParB_F_^1-42^-mCherry-EcoRI^E111Q^. The parameters of ATPase stimulation curves in these and subsequent figures are summarized in Table 2. **Source Data 1:** Numerical data for figure 2 panels A-E (Microsoft Excel). **Source Data 2:** Numerical data for figure 2 panels F, G (Microsoft Excel). **Figure supplement 1:** *G*el filtration column elution profile of ParB_F_^1-42^-mCherry. **Figure supplement 2—Source data 1:** FRAP of ParA_F_-eGFP and ParB_F_^1-42^-mCherry on DNA-carpet. **Figure supplement 3:** Comparison of possible structural domain arrangements of artificially dimeric ParB_F_^1-42^-mCherry-EcoRI^E111Q^ to ParB_F_ structures in the absence or presence of C-nucleotide. **Figure supplement 4—Source data 1:** ParA_F_ ATPase stimulation by ParB_F_^1-42^-mCherry-EcoRI^E111Q^ is not influenced by the addition of DNA fragment containing EcoRI recognition sequence.

**Figure 2—figure supplement 1.**
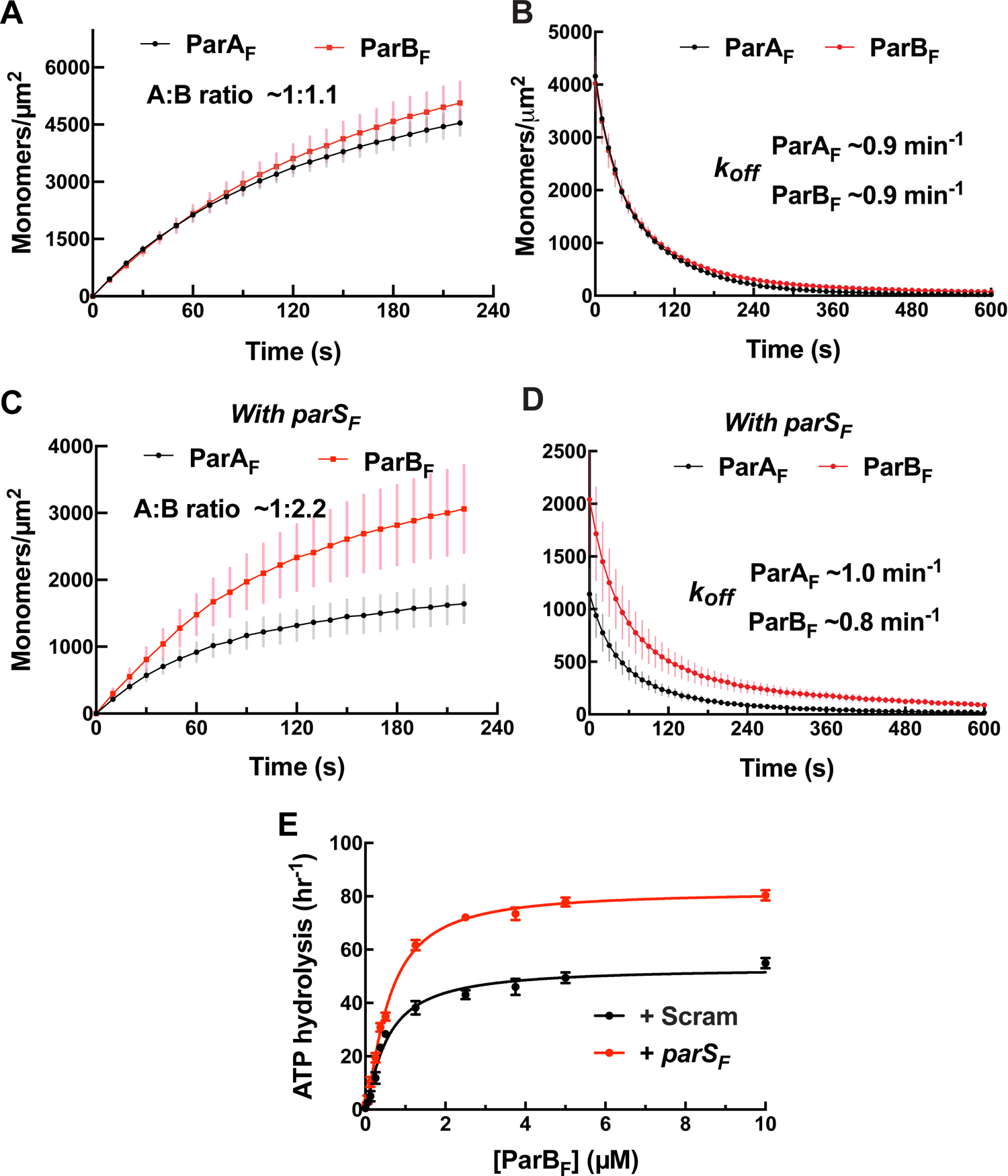
*G*el filtration column elution profile of ParB_F_^1-42^-mCherry. The elution profile of ParB_F_^1-42^-mCherry on a Superose 6 3.2/300 column (GE Healthcare) with a standard curve generated using a protein standards kit (GE Healthcare). The molecular weight of ParB_F_^1-42^-mCherry was estimated to be ∼35 kDa, close to the predicted molecular weight for a monomer of ∼33 kDa.

**Figure 2—figure supplement 2.**
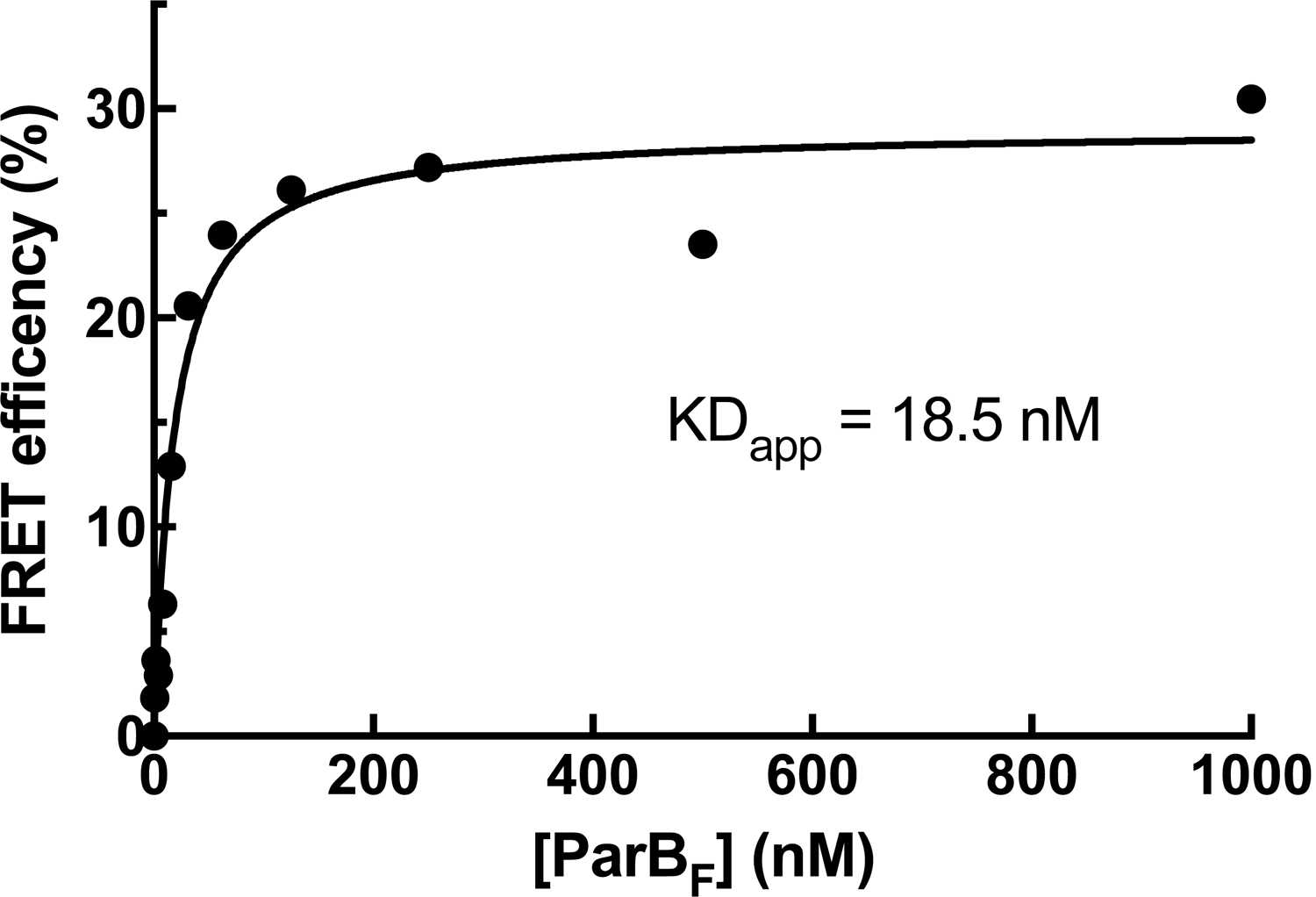
FRAP of ParA_F_-eGFP and ParB_F_^1-42^-mCherry on DNA-carpet. ParB_F_^1-42^-mCherry exchanged significantly faster than ParA_F_ after photo-bleaching. Data have been normalized to the maximum fluorescence recovery. The data were fitted to a single-exponential curve for ParA_F_ and a double-exponential curve for ParB_F_, which was not well fit with a single-exponential recovery curve. ParA_F_ recovered with k_excg_ ∼2.0 min^-1^, and ParB_F_ recovered with k_excg_ ∼15 min^-1^ for the faster (68%), and ∼2.0 min^-1^ for the slower, recovery components. Multi-exponential recovery kinetics can be explained by the relatively large bleached area causing variable diffusion distance for the unbleached molecules need to travel for full fluorescence recovery of the bleached area (this affects recovery of faster recovering components more significantly). An alternative explanation that assumes a 30% population of stable ParA_F_-ParB_F_^1-42^-mCherry complex is unlikely since the buffer wash experiment (Figure 2D) did not support the presence of such a species.

**Figure 2—figure supplement 3.**
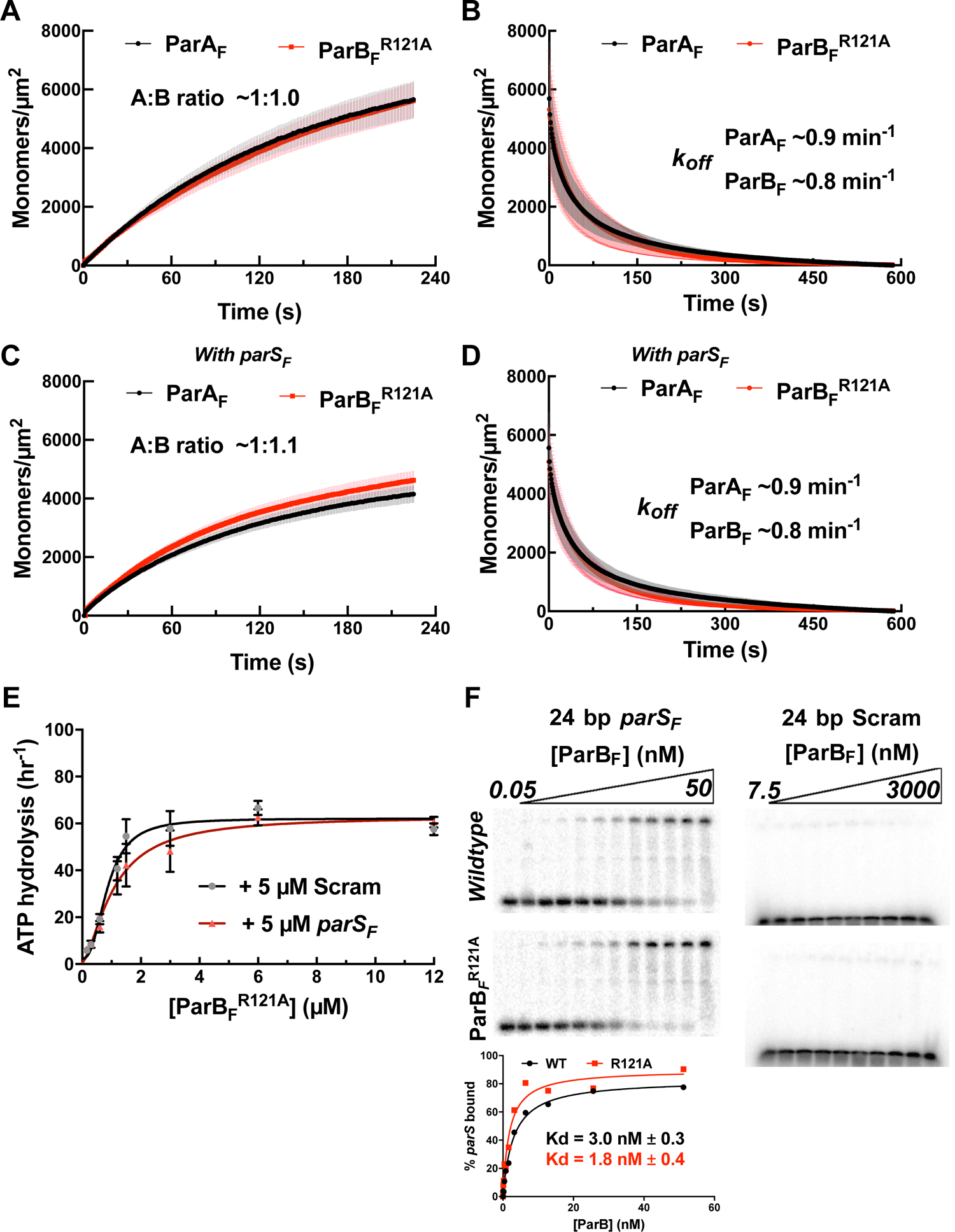
Comparison of possible structural domain arrangements of artificially dimeric ParB_F_^1-42^-mCherry-EcoRI^E111Q^ (A), ParB_F_ dimer structure in the absence (B) and presence (C) of C-nucleotide. This figure depicts montages of domain structures connected by gray lines representing presumed flexible linkers. The structures of the protein domains are based on the following PDB files: EcoR1 (dark green): 1qc9, mCherry (pale yellow): 2h5q, ParB dimerization domain (gray): 1zx4, ParB DNA-binding domain (dark blue) plus CTPase domain (pale blue) in the absence of C-nucleotide: 4umk, ParB DNA-binding domain plus CTPase domain bound to CDP: 6sdk. The dark red helix symbolizes the ParA_F_ interaction domain, which is expected to form an *α*-helix upon binding ParA_F_. The domain arrangement in B is loosely based on the proposal by Chen et al., (2015), which is based on the SAXS envelope of ParB_Hp_ dimer in the absence of C-nucleotide.

**Figure 2—figure supplement 4.**
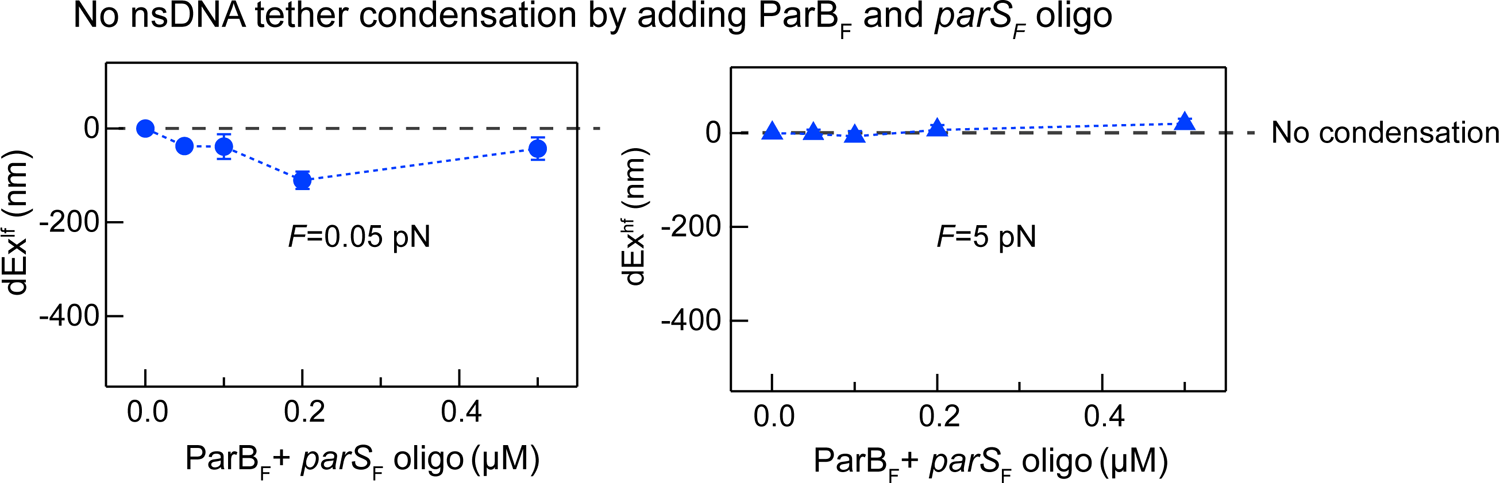
ParA_F_ ATPase stimulation by ParB_F_^1-42^-mCherry-EcoRI^E111Q^ is not influenced by the addition of DNA fragment containing EcoRI recognition sequence. (A) ParA_F_-ATPase activation curve of ParB_F_^1-42^-mCherry-EcoRI^E111Q^ in the absence of the EcoRI sequence duplex (Figure 2G) is shown for comparison. (B) ParA_F_-ATPase activity was measured in the presence of EcoRI-digested pBR322 DNA (60 µg/ml), 77 bp duplex DNA fragment containing two EcoRI recognition sequences (5’-GAATTCCGAGTGGGACCGTGGTCCCAGTCTGATTATCAGACCGAGAATTCAAGTTGGGACCGTGGTCCCAAGAGAAT, plus complement; 1.5 µM) and different concentrations of dimeric ParB_F_^1-42^-mCherry-EcoRI^E111Q^.

**Table 1.**
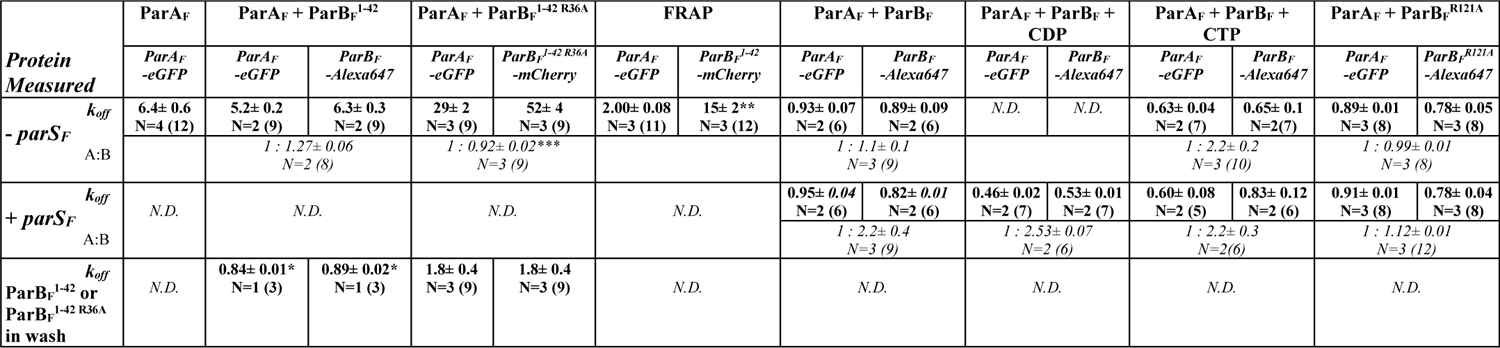
Apparent disassembly or exchange rate constants (min^-1^) and ParB_F_/ParA_F_ ratio from fits of TIRFM wash and FRAP experiments. The apparent dissociation (or FRAP) rate constants (*k_off_*) were obtained for individual time-trajectories by single-exponential curve fitting (except ** where the rate of the faster decay, (68 +/- 0.5%) of a double-exponential fit is shown), and the mean and SEM for the set of independent experiments are shown in **bold** (except *where standard deviation among non-independent repeats within an experiment is shown). (N is the number of separate experiments, with total number of binding/wash cycles for repeated data collection in parenthesis.) ParA:ParB ratios were calculated from the final phase of the individual association time-trajectories (except *** where it was based on the beginning part of the washing phase in the presence of ParB_F_^1-42 R36A^), and the mean and SEM for the set of independent experiments are shown in *italics*. N.D., not done.

**Table 2.** ATPase fit parameters. ATPase measurements were performed with ParA_F_ (1 μM) and different mutants of ParB_F_, 60 μg/ml EcoRI-digested pBR322 DNA plus Scram- or *parS_F_*-DNA fragment and CTP or CDP, as indicated in the column headings. Assays were repeated “*N*” times, each data set of an assay was fit after subtraction of background measured without ParA_F_ to a modified Hill equation: v - v_0_ = (v_max_ [B]^n^) / (K_A_^n^ + [B]^n^), and the mean and standard error of the mean (SEM) of the fit parameters for the *N* measurements are shown. For [B] on the x-axis, total ParB_F_ concentration was used instead of free ParB_F_ concentration due to technical issues in estimating the free ParB_F_ concentration and the meanings of **K_A_** and the cooperativity factor **n** here differ from those in the standard adaptation of the Hill equation. ***v_max_*** is the maximum stimulated ParA_F_ ATPase turnover rate, **K_A_** is the apparent <u>total</u> concentration of ParB_F_ necessary for half maximum stimulation, and **n** is the apparent cooperativity coefficient.

|  | ParB<sub>F</sub> |  | ParB<sub>F</sub> CTP |  | ParB<sub>F</sub> CDP | ParB<sub>F</sub><sup>R121A</sup> |  | ParB<sub>F</sub><sup>1-42</sup> | ParB<sub>F</sub><sup>1-42 R36A</sup> | ParB<sub>F</sub><sup>1-42</sup>-mCherry-EcoRI<sup>E111Q</sup> |  |
| --- | --- | --- | --- | --- | --- | --- | --- | --- | --- | --- | --- |
|  | *Scram* | *parS<sub>F</sub>* | *Scram* | *parS<sub>F</sub>* | *parS<sub>F</sub>* | *Scram* | *parS<sub>F</sub>* |  |  | *Scram* | *EcoRI DNA* |
|  | N = 6 | N = 6 | N = 3 | N = 3 | N = 3 | N = 3 | N = 3 | N = 6 | N = 3 | N = 3 | N = 2 |
| *v<sub>max</sub>* (hr<sup>-1</sup>) | 54<br>± 5 | 79<br>± 2 | 79<br>± 8 | 78<br>± 8 | 87<br>± 5 | 60<br>± 5 | 62<br>± 5 | 83<br>± 3 | 38<br>± 2 | 131<br>± 13 | 125<br>± 9 |
| *K<sub>A</sub>* (μM) | 0.53<br>± 0.09 | 0.59<br>± 0.03 | 0.41<br>± 0.05 | 0.24<br>± 0.02 | 0.34<br>± 0.04 | 0.86<br>± 0.1 | 1.1<br>± 0.1 | 1.2<br>± 0.1 | 108<br>± 13 | 0.16<br>± 0.04 | 0.16<br>± 0.03 |
| Cooperativity coefficient (n) | 1.2<br>± 0.2 | 1.4<br>± 0.1 | 3.3<br>± 1.0 | 3.6<br>± 0.2 | 1.1<br>± 0.1 | 2.5<br>± 0.6 | 1.6<br>± 0.3 | 2.5<br>± 0.3 | 1.5<br>± 0.2 | 1.4<br>± 0.3 | 1.4<br>± 0.2 |

### The N-terminus of ParB_F_ alone enhances DNA binding activity of ParA_F_

We first examined if the N-terminal ParA_F_-activation domain of ParB_F_ (ParB_F_^1-42^) alone can induce the ParA_F_ conformational changes necessary for DNA binding in the presence of ATPγS. The activation domain includes Arginine 36, which is critical for activation of ParA_F_ ATPase (Ah-Seng et al., 2009; Leonard et al., 2005), but lacks the CTPase-(AA63-155), *parS_F_*-binding-(AA160-272), and dimerization-(AA276-323) domains. For these experiments we used ParB_F_^1-42^ fused to the N-terminus of mCherry (ParB_F_^1-42^-mCherry). This protein is a monomer in solution as judged by its elution profile on a gel filtration column (Figure 2—figure supplement 1). ParB_F_^1-42^-mCherry (10 μM) and ParA_F_-eGFP (1 μM) bound the DNA-carpet together in the presence of ATPγS to a density of ∼5000 monomers/μm^2^ maintaining ∼1:1 stoichiometry (Figure 2C). When washed with a buffer containing ATPγS, ParB_F_^1-42^-mCherry dissociated first, with an apparent dissociation rate constant of ∼5.7 min^-1^, closely followed (within a few seconds) by ParA_F_-eGFP dissociation (Figure 2D). On the other hand, when the wash buffer contained 10 μM ParB_F_^1-42^-mCherry and ATP*γ*S, both proteins dissociated together, significantly slower than ParA_F_-eGFP bound to the DNA-carpet alone, with an apparent rate constant of ∼0.9 min^-1^, maintaining ∼1:1 stoichiometry (Figure 2E). After essentially complete dissociation of ParA_F_-eGFP from the DNA-carpet, 10 μM ParB_F_^1-42^-mCherry present in the wash solution showed no significant binding to the DNA-carpet, indicating low intrinsic affinity of this protein for DNA. FRAP measurements of ParA_F_-eGFP— ParB_F_^1-42^-mCherry bound in steady state to the DNA-carpet in the presence of ATPγS also indicated rapid exchange of ParB_F_^1-42^-mCherry (Figure 2—figure supplement 2). Thus, at saturating ParB_F_^1-42^-mCherry concentration, a ParA_F_—ATPγS dimer is bound by two molecules of ParB_F_^1-42^-mCherry occupying both sides of the ParA_F_ dimer. The data also indicate that ParA_F_ dimers adopt a state of slowed dissociation from nsDNA when both of the ParB_F_-interacting faces are occupied by the ParB_F_ N-terminal domain. The nsDNA dissociation rate constants of ParA_F_— ParB_F_ complexes (including those involving ParB_F_ variants) and ParB_F_:ParA_F_ stoichiometry reported above and in the following sections are summarized in Table 1.

### ParA_F_ ATPase activation requires binding of two copies of ParB_F_ N-terminal domain to the ParA_F_ dimer

ParB_F_^1-42^ stimulated ParA_F_-ATPase (1 μM) with a clear sigmoidal ParB_F_^1-42^ concentration dependence and a half-maximum activation concentration of ∼1.2 μM (Figure 2F; Table 2). Thus, monomeric ParB_F_^1-42^ appears to activate ParA_F_-ATPase when it binds on both sides of the DNA-bound ParA_F_ dimers. To test if the observed sigmoidal concentration dependence is due to the monomeric nature of ParB_F_^1-42^, we prepared the N-terminal region of ParB_F_ fused to mCherry and the nuclease activity deficient EcoRI^E111Q^, ParB_F_^1-42^-mCherry-EcoRI^E111Q^ (see Figure—2 figure supplement 3A). This construct, with expected EcoRI dimerization *K_D_* < 20 pM (Modrich and Zabel, 1976), efficiently activated ParA_F_-ATPase at least to a similar maximum rate as ParB_F_^1-42^, but with a K_half_ of ∼0.15 μM, ∼8-fold lower than ParB_F_^1-42^, and displayed no noticeable sigmoidal concentration dependence (Figure 2G). Potential binding of the inactive EcoRI domain to DNA did not appear to influence the ATPase activation properties of this construct; addition of EcoRI-binding DNA fragment in the reaction did not impact the ATPase activation (Figure 2—figure supplement 4). Based on these results, we conclude that both ParB_F_-binding faces of a ParA_F_ dimer must be occupied by ParB_F_ N-termini for stimulation of its ATPase activity.

### ParB_F_^1-42 R36A^ forms a rapidly disassembling complex with ParA_F_ on the DNA carpet

An R36A mutation was reported to significantly compromise ParB_F_’s ability to activate ParA_F_’s ATPase (Ah-Seng et al., 2009). To test whether this mutation affected ParB_F_’s ability to form a complex with ParA_F_ we repeated the experiments shown in Figure 2C-E using ParB_F_^1-42 R36A^-mCherry. ParB_F_^1-42R36A^-mCherry and ParA_F_-eGFP bound with an approximately 1:1 stoichiometry, similar to ParB_F_^1-42^-mCherry but reached a steady-state density on the DNA-carpet of less than 10% of the density observed with ParB_F_^1-42^-mCherry (Figure 3A). When washed with buffer containing ATP*γ*S, ParB_F_^1-42 R36A^-mCherry disassociated first followed by ParA_F_, similar to the results obtained with ParB_F_^1-42^-mCherry but ParB_F_^1-42 R36A^-mCherry dissociated ∼10-fold faster, followed by dissociation of ParA_F_-eGFP within a few seconds (Figure 3B). When the wash buffer also contained 10 μM ParB_F_^1-42 R36A^-mCherry the two proteins dissociated in parallel maintaining ∼1:1 stoichiometry (Figure 3C). Together these observations indicate that ParB_F_^R36A^ interacts with ParA_F_, but with a much faster dissociation rate constant compared to wild-type ParB_F_. ParB_F_^1-42 R36A^ could activate ParA_F_-ATPase with an increased half-saturation concentration of 108 µM, approximately 100-fold higher than ParB_F_^1-42^ (Figure 3D). These results explain the puzzling report that while the R36A mutation severely compromised activation of ParA_F_-ATPase by ParB_F_, it did not impede oscillation of ParA_F_ on the nucleoid, and only mildly reduced plasmid stability (Ah-Seng et al., 2013). At the interface between the ParA_F_-bound nucleoid and partition complexes containing many ParB_F_ dimers, the local ParB_F_ concentration is expected to be sufficiently high for this mutant protein to activate ParA_F_-ATPase to effectively generate a ParA_F_ depletion zone and motive force driving the partition complex as indicated by the repeated oscillation of the nucleoid-bound ParA_F_ distribution.

**Figure 3.**
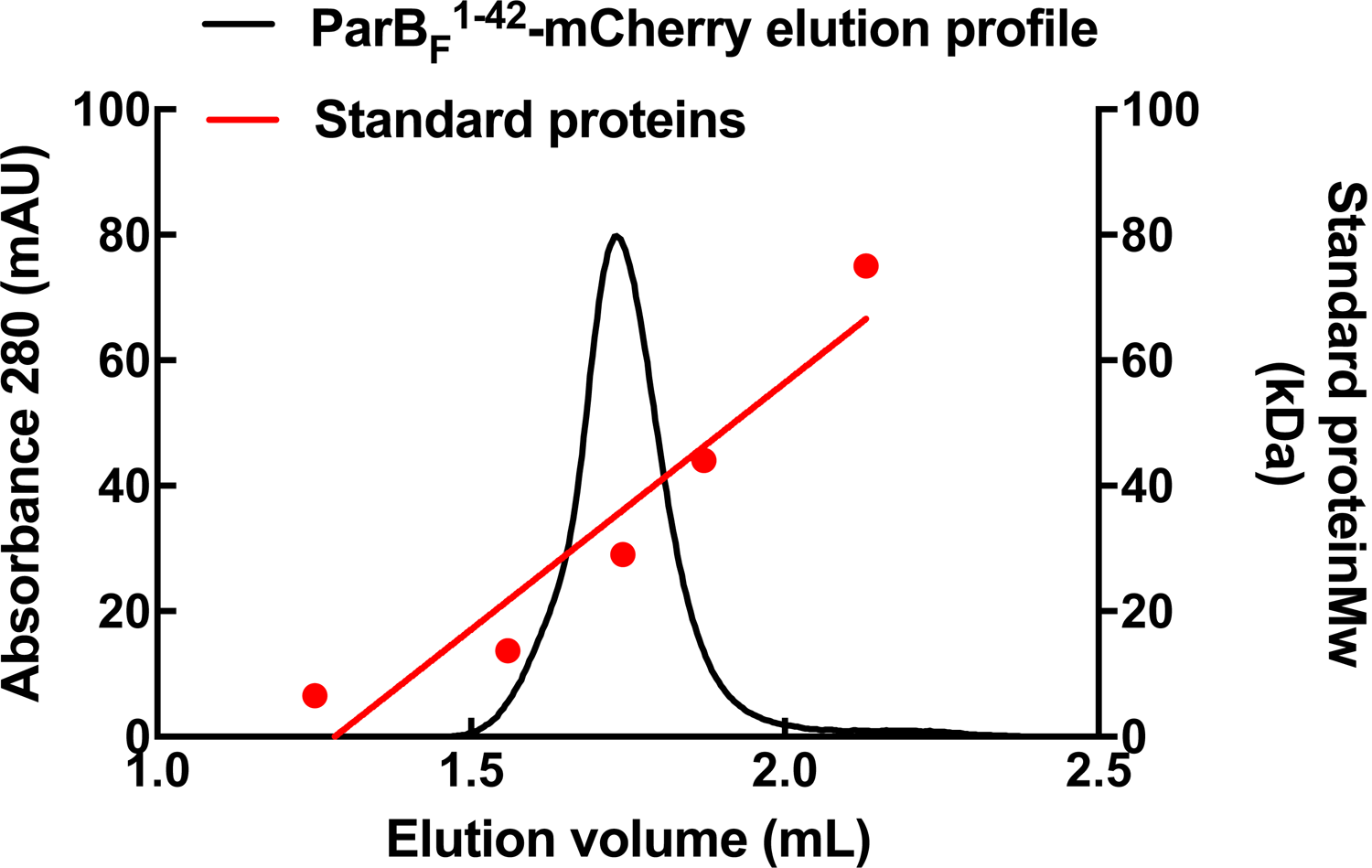
ParB_F_^1-42 R36A^-mCherry dissociates faster from nsDNA-carpet-bound ParA_F_-ATPγS dimer, and ParA_F_-ATPase activation requires higher ParB_F_^1-42 R36A^ concentration. (A) ParB_F_^1-42 R36A^-mCherry (10 μM) and ParA_F_-eGFP (1 μM) preincubated with ATPγS were infused into the nsDNA-carpeted flow cell and then (B) washed with buffer containing ATPγS. (C) The washing experiment of B was repeated with buffer containing ATPγS and ParB_F_^1-42 R36A^-mCherry (10 μM). (D) ParA_F_-ATPase activity was measured in the presence of EcoRI-digested pBR322 DNA (60 μg/ml) as a function of ParB_F_^1-42 R36A^ concentration. See Figure 2 legend and Tables 1 and 2 for additional details. **Source data 1:** Numerical data for figure 3 panels A-C (Microsoft Excel). **Source data 2:** Numerical data for figure 3 panels D (Microsoft Excel).

### ParA_F_ and ParB_F_ bind to and dissociate from nsDNA together in the presence of ATPγS with ∼1:1 stoichiometry

When ParA_F_-eGFP and full-length ParB_F_-Alexa647 were incubated together at 1 µM and 2 µM, respectively in the presence of ATPγS, they bound to the DNA-carpet in parallel maintaining ∼1:1 stoichiometry up to a density of ∼5000 monomers/μm^2^ (Figure 4A). They also dissociated from the DNA-carpet in parallel, maintaining ∼1:1 stoichiometry, when washed with a buffer containing ATPγS, with an apparent dissociation rate constant of approximately ∼1 min^-1^ (Figure 4B). These results show that ParA_F_ and ParB_F_ form a hetero-tetramer containing two monomers each of ParA_F_ and ParB_F_ (A_2_B_2_), or larger oligomers composed of the hetero-tetramers, that binds as a unit on nsDNA in the presence of ATPγS.

**Figure 4.**
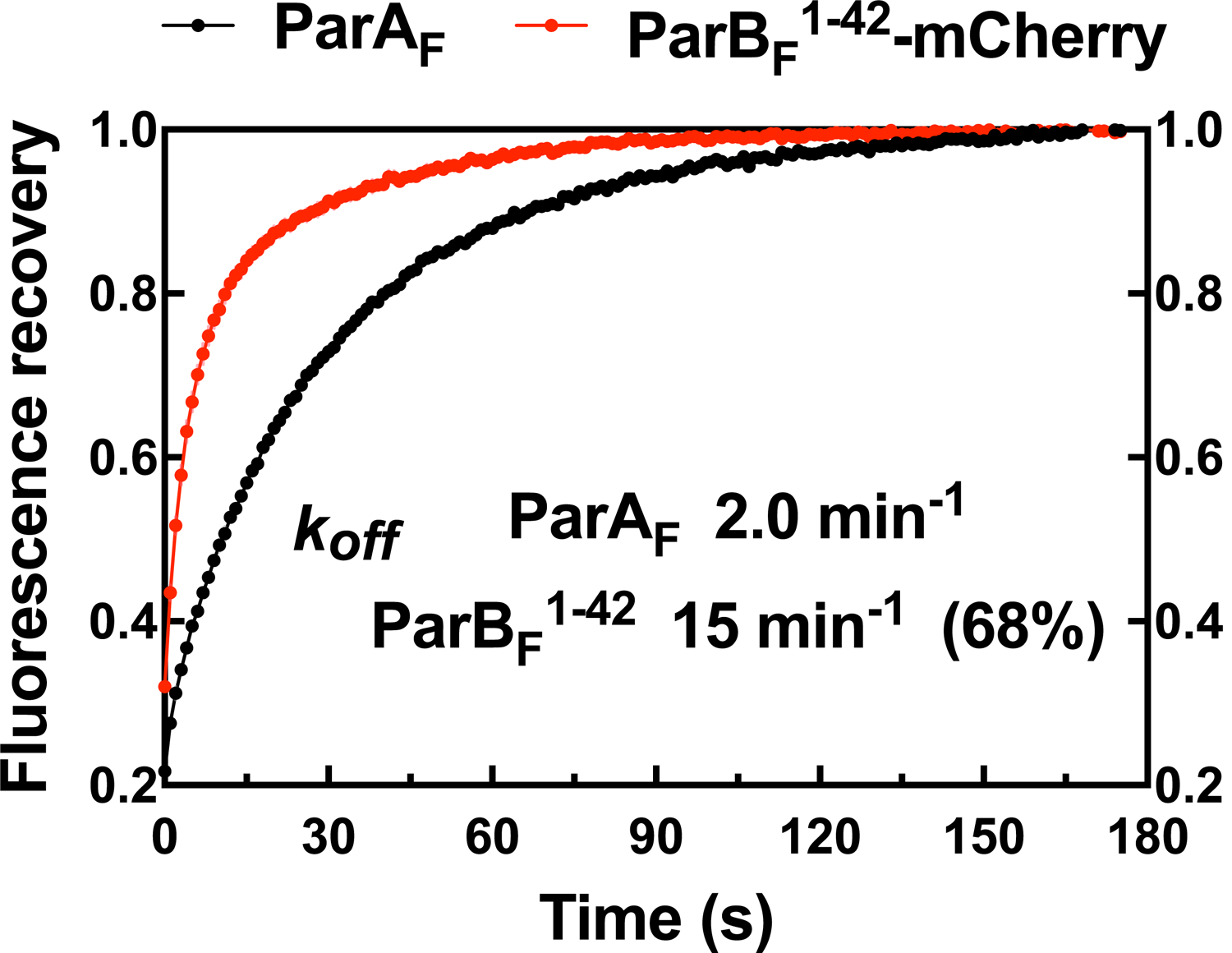
parS_F_ DNA alters protein stoichiometry of the ParA_F_—ParB_F_ complex formed prior to ATP hydrolysis and the extent of ParA_F_-ATPase activation by ParB_F._ (A) ParA_F_-eGFP (1 μM) and ParB_F_-Alexa647 (2 μM) preincubated with ATPγS were infused into the nsDNA-carpeted flow cell and then (B) washed with buffer containing ATPγS. (C and D) As A and B except the sample included 24bp *parS_F_* DNA fragment (1.1 μM). (E) ParA_F_-ATPase activity was measured in the presence of EcoRI-digested pBR322 DNA (60 μg/ml), different concentrations of ParB_F_ and either a *parS_F_*-DNA fragment or a DNA fragment with a scrambled sequence (1.1-fold higher concentrations than the ParB_F_ dimers). See Figure 2 legend and Tables 1 and 2 for additional details. **Source data 1:** Numerical data for figure 4 panels A-D (Microsoft Excel). **Source data 2:** Numerical data for figure 4 panels E (Microsoft Excel). **Figure supplement 1—Source data 1:** Numerical data for Figure 4—figure supplement 1 (Microsoft Excel). **Figure supplement 2—Source data 1:** Numerical data for Figure 4—figure supplement 2 (Microsoft Excel).

**Figure 4—figure supplement 1.**
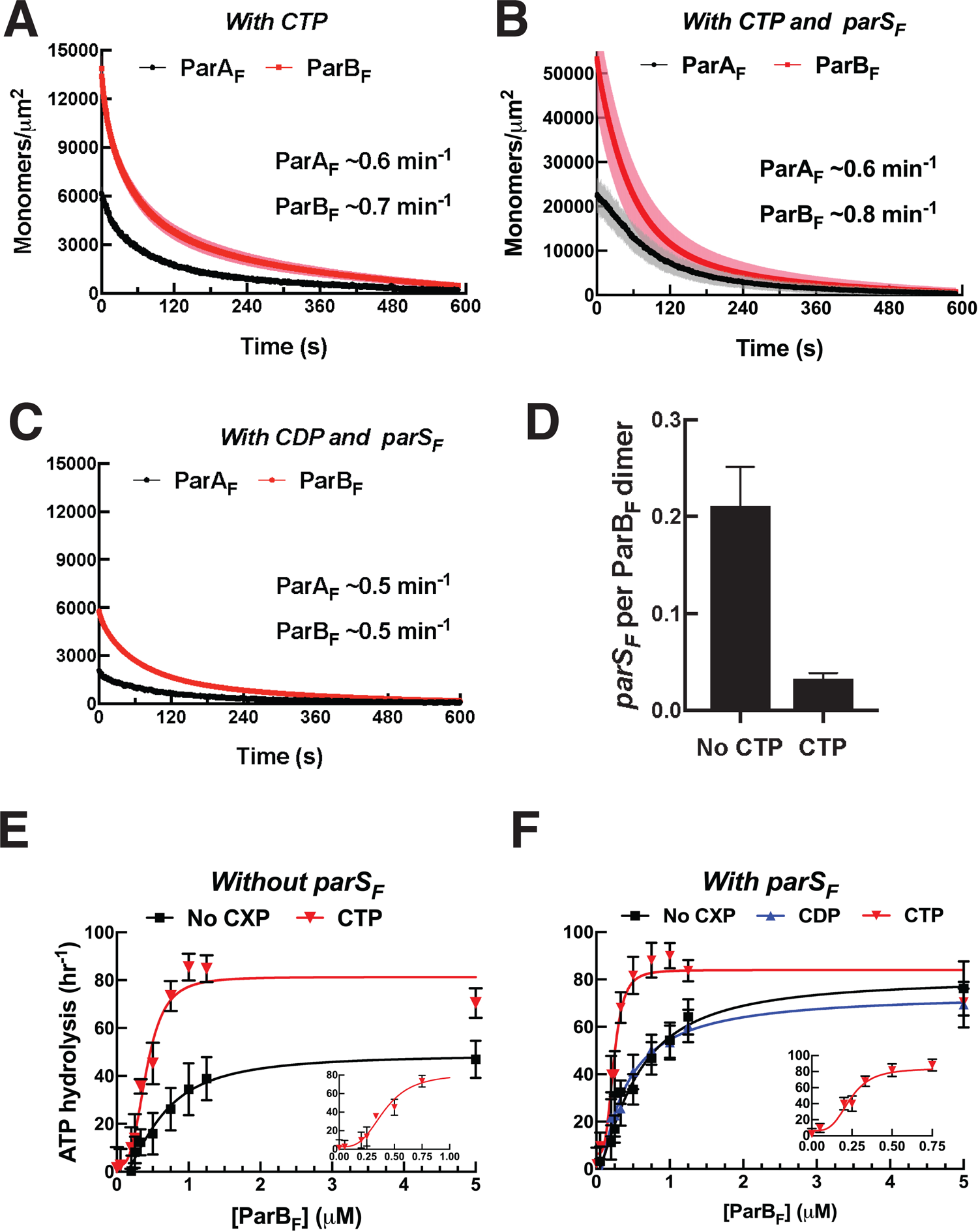
Determination of ParB_F_ monomer-dimer *K_D_* by FRET. In order to estimate the *K_D_* for dimerization of ParB_F_ two batches of protein were separately labeled with Alexa 488 or Alexa 594 then mixed at equal concentrations, the sum of which are shown on the *x*-axis. The FRET efficiency was measured using a fluorescence plate reader (Clariostar Plus, BMG Labtech). Assuming the observed FRET efficiency linearly correlates to the fraction of the protein in the dimer state, the apparent *K_D_* for dimerization of ParB_F_ was estimated to be ∼18.5 nM.

**Figure 4—figure supplement 2.**
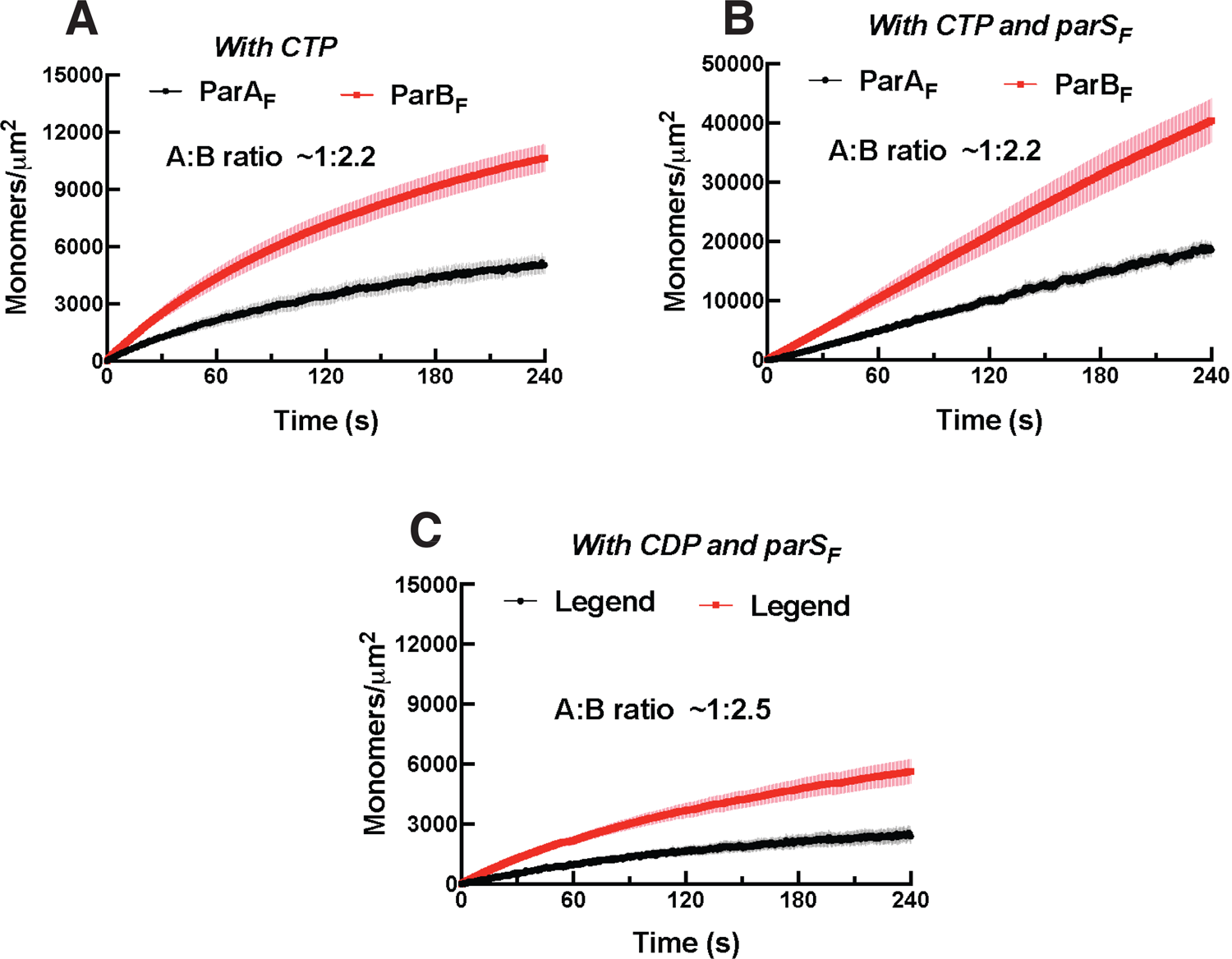
Mutation of a BoxII residue R121A does not affect the affinity of ParB_F_^R121A^ for *parS_F_*, but neither the protein stoichiometry of the ParA_F_-ParB_F_^R121A^ complex assembled on nsDNA prior to ATP hydrolysis, nor the extent of ParA_F_-ATPase activation by ParB_F_^R121A^ is impacted by the presence of *parS_F_*. ParA_F_ (1 μM) and ParB_F_ ^R121A^ (2 μM) were preincubated in the presence of ATPγS either without (A) or together with 24 bp *parS_F_* DNA fragment (1.1 μM, C), infused into to the DNA-carpeted flow cell and washed with buffer containing ATPγS (B, D). (E) ParA_F_ ATPase activation by ParB_F_^R121A^ was measured in the presence of EcoRI-digested pBR322 DNA (60 μg/ml) and different concentrations of ParB_F_^R121A^ either in the absence (black line) or presence (red line) of *parS_F_* (at 10% excess of the dimer concentrations of ParB_F_^R121A^). (F) The binding of ParB_F_ or ParB_F_^R121A^ to 0.1 nM P^32^-labeled 24 bp *parS_F_* DNA (left) or to 0.1 nM P^32^-labeled 24 bp *scrambled* DNA (right) was measured by EMSA. A 4-12% gradient acrylamide gel was run in 1x TBE + 5 mM Mg_2_Ac at 4 °C for 1 hour, 150 V. The gels were quantified (bottom) to estimate the *parS* binding *K_D_* of ParB_F_ and ParB_F_^R121A^.

Full-length ParB_F_, which forms dimer with apparent *K_D_* of ∼ 19 nM (Figure 4—figure supplement 1), activated ParA_F_-ATPase in the presence of nsDNA to ∼50 hr^-1^ without significant sigmoidal concentration dependence (Figure 4E). Based on the results of experiments with monomeric ParB_F_^1-42^ proteins described earlier, we conclude that a single dimer of full-length ParB_F_ can straddle an nsDNA-bound ParA_F_ dimer, permitting the two N-termini to interact with both of the ParB_F_-binding faces of the ParA_F_ dimer to activate the ATPase.

### In the presence of parS_F_, ParB_F_ forms a 2:1 complex with ParA_F_

We next asked if ParB_F_ bound to *parS*_F_ interacts differently with ParA_F_ on the DNA-carpet. We preincubated ParA_F_-eGFP, ParB_F_-Alexa647, ATPγS and a 24 bp duplex DNA fragment containing a single *parS*_F_ consensus sequence, at a slight molar excess over ParB_F_ dimer, for 10 min at room temperature. At the concentrations used, most of the ParB_F_ dimers are expected to be bound to *parS*_F_. When infused into the DNA-carpeted flow cell, ParA_F_-eGFP and ParB_F_-Alexa647 bound to and dissociated from the carpet with a stoichiometry of ∼1:2 (Figure 4C, D), in sharp contrast to the ∼1:1 stoichiometry without *parS*_F_ DNA. The kinetic parameters of the complex assembly and disassembly were not significantly affected. These results demonstrate that ParA_F_ and ParB_F_ form a complex of one ParA_F_ dimer and two ParB_F_ dimers (A_2_B_4_) in the presence of *parS_F_*.

Does the change in protein stoichiometry caused by *parS_F_* translate to different levels of ParA_F_-ATPase activation? A previous study, comparing plasmid DNA with and without *parS_F_* as the cofactor, showed that ParB_F_ activates ParA_F_-ATPase a few-fold more efficiently in the presence of plasmid DNA containing a full *parS_F_* site (Ah-Seng et al., 2009). We titrated ParB_F_ in the presence of ParA_F_, pBR322 DNA and 24 bp *parS_F_* duplex at a stoichiometric excess concentration over the ParB_F_ dimer. In the presence of *parS_F_* DNA, ParB_F_ activated ParA_F_-ATPase to a maximum turnover rate of ∼80 hr^-1^, a ∼60% increase compared to reactions where the *parS_F_* fragment was replaced with a scrambled sequence fragment (Figure 4E). These results indicate that a single *parS*_F_ DNA-bound ParB_F_ dimer cannot straddle an nsDNA-bound ParA_F_ dimer to activate the ATPase, but by binding two ParB_F_ dimers the ATPase activation level reaches slightly higher level than in the absence of *parS*_F_ DNA.

We note that ParB_F_^R121A^, harboring a mutation in the conserved Box II region of the CTPase domain, neither exhibited a change in the ParB_F_/ParA_F_ complex stoichiometry, nor a change in the ParB_F_-stimulated ATP turnover, in response to *parS_F_* (Figure 4—figure supplement 2), suggesting that the effects of *parS_F_* binding described above are mediated through conformational changes in the CTPase domain (see below for further discussion).

### CTP alters the complex formed between ParA_F_ and ParB_F_ in a manner similar to parS_F_, and accelerates complex formation in the presence of parS_F_

ParB proteins have recently been reported to have CTPase activity that is coupled to changes in their DNA binding properties and refolding of the CTPase domains into a globular dimeric structure in the presence of CTP (Soh et al., 2019; Osorio-Valeriano et al., 2019) from the more extended and poly-dispersed structure in the absence of nucleotide (Chen et al., 2015). We therefore decided to test if the addition of CTP influences the ParA_F_—ParB_F_ complex formed on the DNA-carpet in the presence of ATPγS. When ParA_F_-eGFP and ParB_F_-Alexa647 were incubated together in the presence of ATPγS (1 mM) and CTP (2 mM), they bound to and dissociated from the nsDNA-carpet with a stoichiometry of ∼1:2 (Figure 5A, Figure 5—figure supplement 1A). The assembly kinetics of the carpet-bound complex was roughly the same as in the absence of CTP, however the apparent dissociation rate constant during buffer wash was slightly but reproducibly slower by a factor of roughly two at ∼ 0.6 min^-1^. When *parS_F_* was included together with CTP, the rate of A_2_B_4_ complex assembly on the DNA-carpet increased several-fold, the binding density of the complex on the DNA-carpet reached a correspondingly higher level, and the two proteins dissociated from the DNA-carpet maintaining a ∼1:2 stoichiometry with apparent dissociation rate constant similar to that in the absence of *parS_F_* (Figure 5B, Figure 5—figure supplement 1B). When CTP was replaced by CDP in the presence of *parS_F_*, although the ParB_F_/ParA_F_ ratio remained above 2, unlike in the presence of CTP the complex assembly rate did not increase (Figure 5C, Figure 5—figure supplement 1C), thus behaving similarly to the reaction in the presence of *parS_F_* alone.

**Figure 5.**
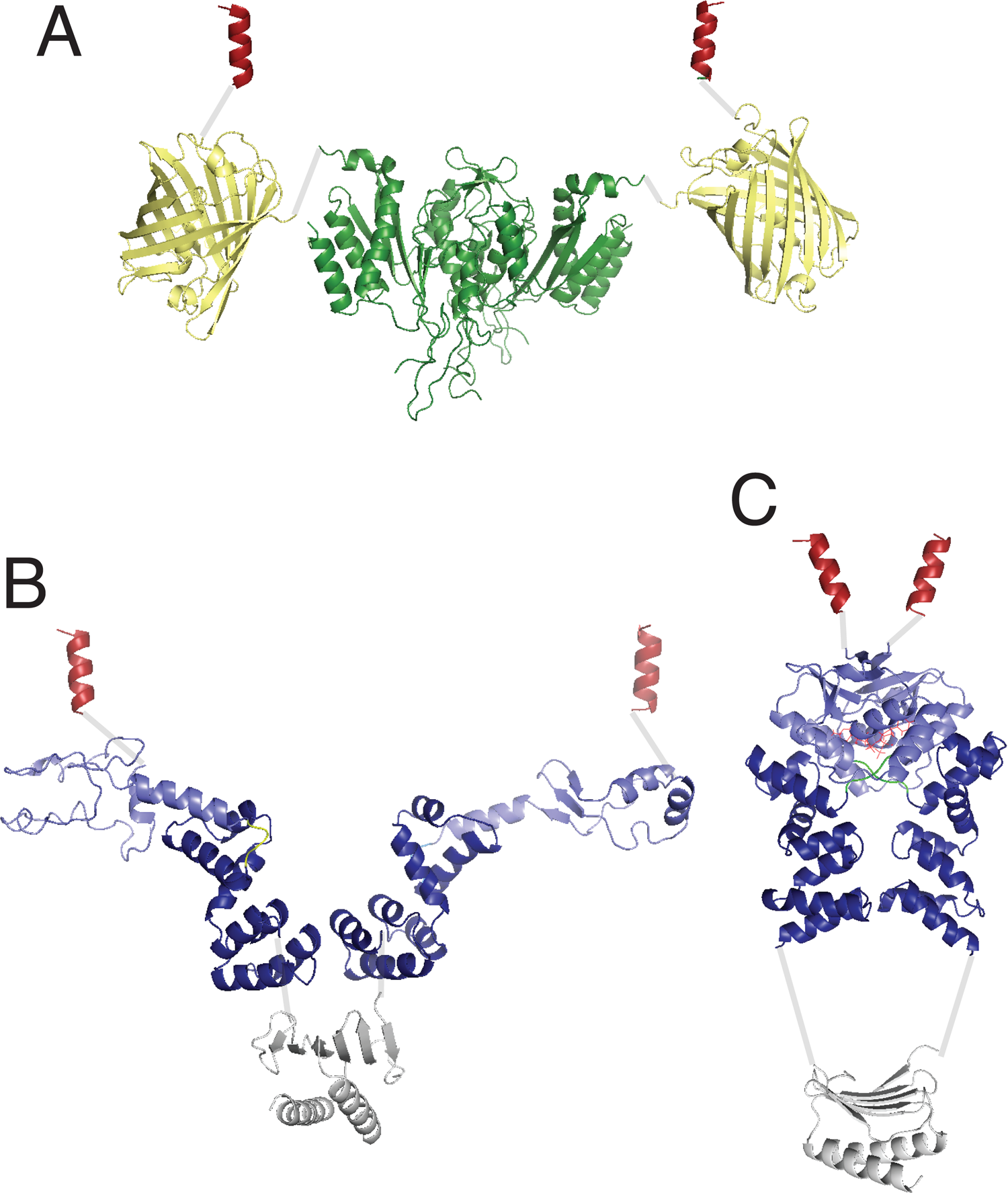
*CTP and parS_F_ together alter interactions between ParB_F_ and ParA_F_ dimers.* (A) ParA_F_-eGFP (1 μM) and ParB_F_-Alexa647 (2 μM) preincubated with ATPγS and CTP (2mM) were infused into the nsDNA-carpeted flow cell and then washed with buffer containing ATPγS and CTP. (B) As in A, except a 24 bp *parS_F_* fragment (1.1 μM) was added to the sample mixture. (C) As in B, except CTP was replaced by CDP. For binding curves, see Figure 5—figure supplement 1A-C. (D) ParA_F_ (1 μM), ParB_F_-Alexa647 (2 μM) and Alexa488-labeled 24 bp *parS_F_* fragment (1.1 μM) preincubated with ATPγS or ATP*γ*S plus CTP (2mM) were infused into the nsDNA-carpeted flow cell and after 240 sec, the ratio of the carpet-bound *parS_F_* fragment and ParB_F_ dimer was measured. (E) ParA_F_-ATPase activity was measured in the presence of EcoRI-digested pBR322 DNA (60 μg/ml), different concentrations of ParB_F_ and either no C-nucleotide, or 2mM CTP. Inset shows data in the presence of CTP with expanded abscissa. (F) As in E except the reactions also contained 24 bp *parS_F_* fragment (1.1-fold higher concentrations than ParB_F_ dimers). Inset shows data in the presence of *parS_F_* and CTP with expanded abscissa. See Figure 2 legend and Tables 1 and 2 for additional details. **Source data 1:** Numerical data for figure 5 panels A-D (Microsoft Excel). **Source data 2:** Numerical data for figure 5 panels E, F (Microsoft Excel). **Figure supplement 1—Source data 1:** Numerical data for Figure 5—figure supplement 1 (Microsoft Excel). **Figure supplement 2—Source data 1:** Numerical data for Figure 5—figure supplement 2 (Microsoft Excel).

**Figure 5—figure supplement 1.**
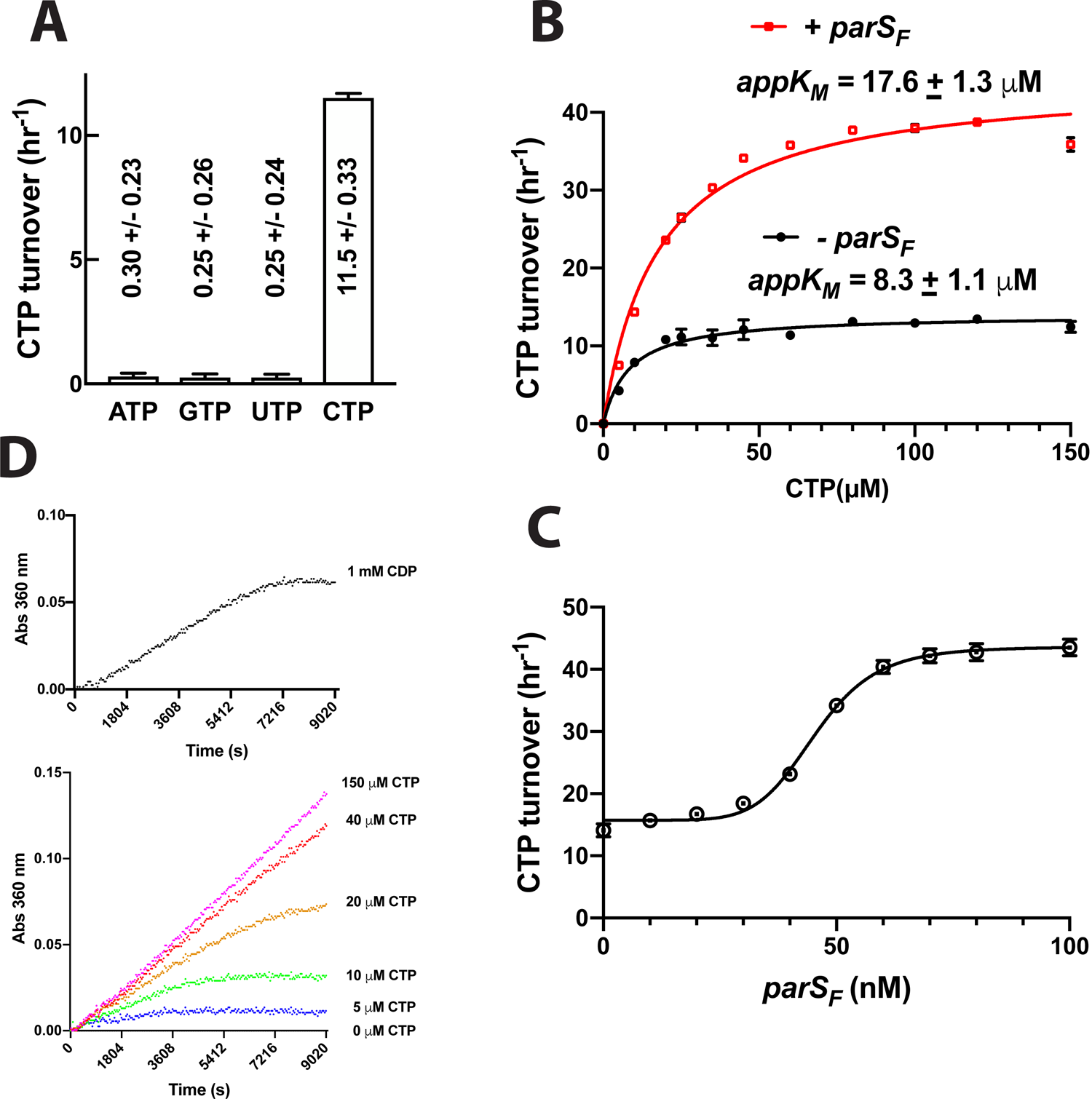
Binding curves for ParA_F_ and ParB_F_ with CDP or CTP associating with the DNA-carpet. (A) ParA_F_-eGFP (1 μM) and ParB_F_-Alexa647 (2 μM) preincubated in the presence of ATPγS and 2 mM CTP were infused into the nsDNA-carpeted flow cell. (B) ParA_F_-eGFP (1 μM) and ParB_F_-Alexa647 (2 μM) preincubated in the presence of 24bp *parS_F_* DNA fragment (1.1 μM), ATPγS and 2 mM CTP were infused into the nsDNA-carpeted flow cell. (C) ParA_F_-eGFP (1 μM) and ParB_F_-Alexa647 (2 μM) preincubated in the presence of 24bp *parS_F_* DNA fragment (1.1 μM), ATPγS and 2 mM CDP were infused into the nsDNA-carpeted flow cell.

**Figure 5—figure supplement 2.**
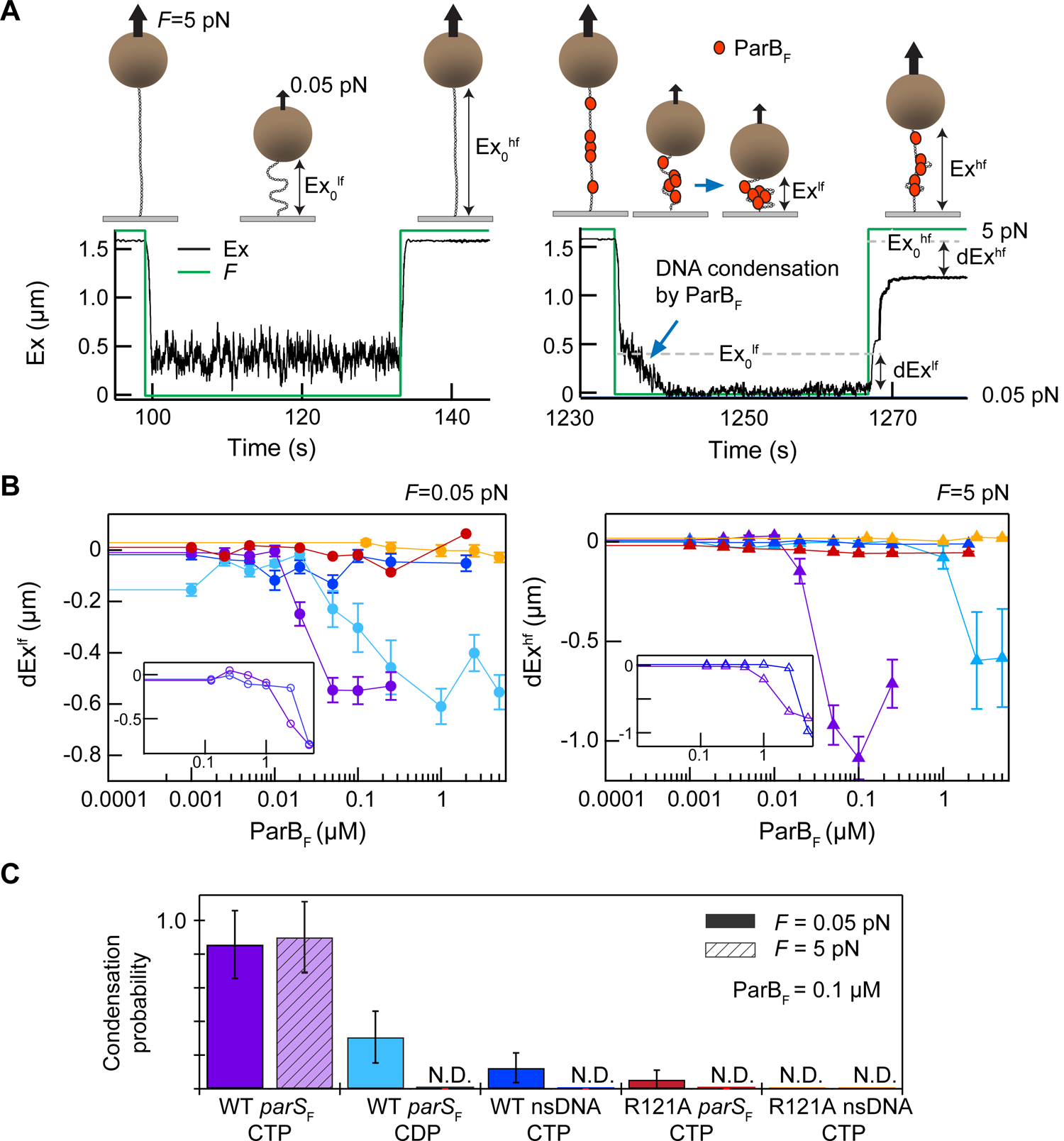
Nucleotide specificity and CTPase activity of ParB_F_ in the presence of different concentrations of CTP, in the presence or absence of *parS_F_*. (A) NTPase activities of ParB_F_ (0.84 μM) in the presence of 100 μM ribonucleoside triphosphates were measured in the CTPase buffer (Materials and Methods). (B) The CTP hydrolysis initial rate by ParB_F_ (0.84 μM) was measured at different CTP concentrations to estimate the apparent CTP *K_M_* with or without 100 nM *parS_F_* (24bp duplex with one copy of ParB_F_ binding consensus sequence). (C) *parS_F_* concentration dependence of the CTPase activity of ParB_F_ (0.84 μM, 100 μM CTP). Pronounced sigmoidal *parS_F_* concentration dependency of the CTPase stimulation was observed. This is in contrast to the hyperbolic *parS_Bsu_* concentration dependency reported for ParB_Bsu_ (Soh et al., 2019). The mechanistic reason for this concentration dependency is currently under investigation. (D) CDP used in this study contained a compound that can be hydrolyzed by ParB_F_ to release Pi. Time course of Pi production from 1 mM CDP in the presence of 0.84 μM ParB_F_ is compared to reactions containing 5 to 150 μM CTP.

Next, we asked if the *parS_F_* DNA fragment was incorporated in the A_2_B_4_ complexes assembled in its presence. The experiments in the presence of *parS_F_* were repeated in the presence or absence of CTP with ParA_F_ (1 μM), ParB_F_-Alexa647 (2 μM) and Alexa488-*parS_F_* (1.1 μM), and the nsDNA-carpet-bound ratio of *parS_F_* and ParB_F_ after 240 sec sample infusion was measured (Figure 5D). The observed *parS_F_* / (ParB_F_)_2_ ratio in the absence of CTP was ∼0.2, while in the presence of CTP, the ratio was only ∼0.04. Thus, whereas the assembly of the A_2_B_4_ complex involving CTP-ParB_F_ was accelerated by *parS_F_*, a very small fraction of the resulting complex contained the *parS_F_*-DNA fragment, indicating that *parS_F_* plays a catalytic role in the activation of CTP-ParB_F_ and accelerated assembly of the A_2_B_4_ complex. This parallels the observation that a much lower concentration of *parS_F_* fully activated the CTPase activity of ParB_F_ (Figure 5—figure supplement 2C) as has also been shown for ParB_Bsu_ (Soh et al., 2019).

### ParB_F_ activates ParA_F_-ATPase to the full extent without parS_F_ in the presence of CTP

The maximum ParB_F_ activation of ParA_F_-ATPase in the presence of CTP, with or without *parS_F_*, was comparable to that of *parS_F_*-bound ParB_F_ in the absence of CTP (Figure 5E, F). The half-saturation concentration of ParB_F_ in the presence of *parS_F_* and CTP was significantly lower than in the absence of CTP (∼0.24 μM *vs* ∼0.6 μM). Combined with the observation of faster assembly of the complex on the DNA-carpet, a likely possibility is that in the presence of CTP and *parS_F_*, the ParB_F_ dimer adopts a unique state that interacts with ParA_F_ dimers with a higher association rate constant. We note that the ParA_F_-ATPase assays in this study measured radioactive *γ*-phosphate release from *γ*-^32^P-ATP, avoiding potential technical complications associated with ATPase measurements in the presence of CTP.

We next measured the ParB_F_-CTPase activity to estimate the apparent *K_M_* and *k_cat_* of ParB_F_ for CTP hydrolysis in the presence and absence of *parS_F_*. ParB_F_ had negligible activity for all NTPs other than CTP (Figure 5—figure supplement 2A), and the CTP hydrolysis rate increased with a hyperbolic CTP concentration dependence, which could be fit with the Michaelis-Menten equation with apparent *K_M_* of ∼8 μM and ∼18 μM and maximum turnover rates of ∼14 h^-1^ and ∼44 h^-1^ in the absence and presence of *parS_F_* DNA, respectively (Figure 5—figure supplement 2B). Thus, 2 mM CTP used in the experiments of Figure 5 should have remained saturating ParB_F_ for the duration of the reaction. Stimulation of the CTPase activity by *parS_F_* exhibited a pronounced sigmoidal concentration dependence approaching saturation at ∼ 60 nM, well below the ParB_F_ concentration in the reaction (0.84 μM) (Figure 5—figure supplement 2C).

During these experiments, which were prompted by a reviewer’s comment, we also attempted to characterize the interaction between CDP and ParB_F_, but discovered that the CDP used here contained ∼2% contamination of a compound that released Pi upon incubation with ParB_F_ (Figure 5—figure supplement 2D). The results shown in Figure 5C and Figure 5F suggest this contamination did not strongly influence the reactions involving CDP, considering that they generally paralleled the results obtained in the absence of C-nucleotides with only minor deviations. However, this contamination prevented us from accurately determining the affinity of ParB_F_ for CDP.

### In the presence of CTP, ParB_F_ condenses DNA carrying parS_F_ in cis

The recently discovered CTP and *parS* dependent ParB conformational change appears to promote ParB *parS*-DNA binding and spreading (Soh et al., 2019), impacting ParB-DNA partition complex assembly. *In vivo*, spreading ParB forms condensed foci around *parS* sites indicating that *parS* driven ParB spreading likely occurs *in cis*. Nonetheless, the possibility that *parS* can trigger ParB spreading *in trans* has not been tested *in vitro*. Previous studies reported DNA condensation by *B. subtilis* ParB *via* ParB—ParB interactions, but these studies were conducted in the absence of CTP and did not observe a strong effect of *parS in cis* (Graham et al., 2014; Song et al., 2017; Taylor et al., 2015). To see if *parS_F_* can act *in-trans* and to characterize how *parS_F_* and CTP influence ParB_F_—DNA interactions *in vitro*, we conducted single-molecule DNA pulling experiments employing magnetic tweezers. ParB_F_ at various concentrations was infused into a flow cell containing ∼ 5 kbp DNA tethers that anchored magnetic beads to the coverslip surface (Figure 6A). The tethers contained either twelve *parS_F_* consensus sequence repeats at their midpoints (*parS_F_*-DNA), or no *parS_F_* sequence (nsDNA). The protein sample was infused while the DNA tethers were stretched at 5 pN force, preventing DNA condensation. To allow DNA condensation by bound ParB_F_ molecules, the force was dropped to 0.05 pN and the tether extension was monitored for 30 sec. To assess the stability of DNA condensation by ParB_F_ dimers, tether extension was monitored after increasing the force to 5 pN. In the absence of CTP, we only observed condensation at very high concentrations of ParB_F_ (>5 µM) and did not see a significant difference between *parS_F_*-containing and non-specific tethers (Figure 6B inset). However, in the presence of CTP, 50 nM ParB_F_ robustly condensed *parS_F_*-containing DNA tethers (Figure 6B purple). These condensed protein-DNA complexes resisted 5 pN extension force, requiring many minutes at 5 pN tension to de-condense (Figure 6—figure supplement 1A). The slow de-condensation took place through a series of abrupt steps, which we interpret as stepwise opening of large DNA loops held by multiple ParB_F_—ParB_F_ interactions (Figure 6—figure supplement 1A). Condensation was comparable for DNA molecules that were topologically constrained, i.e., could be supercoiled, or unconstrained (nicked), suggesting that condensation is not a consequence of topological changes in the DNA caused by ParB_F_ translocating away from parS_F_ sites (Figure 6—figure supplement 2). We observed some condensation events with *parS_F_* containing tethers in the presence of CDP, but these events were rarer, required higher ParB_F_ concentrations, and were almost completely de-condensed within 5 sec of raising the force to 5 pN, in stark contrast to condensation in the presence of CTP (Figure 6B light blue, Figure 6—figure supplement 1B). Since this experiment was also carried out using CDP that contained a compound hydrolysable by ParB_F_, contribution of this compound to the limited tether condensation cannot be ruled out. In contrast, ParB_F_ was unable to substantially condense DNA tethers lacking *parS_F_* sequences, even in the presence of CTP, and rare condensation events were quickly reversed by the application of 5 pN force (Figure 6B dark blue). Addition of *parS_F_*-containing DNA fragments together with ParB_F_ and CTP did not rescue the inability to condense tethers lacking *parS_F_*, indicating that *parS_F_* cannot act *in trans* to promote ParB spreading and condensation of DNA molecules (Figure 6—figure supplement 3). Together these results indicate that *parS_F_* mediates loading of multiple CTP-bound ParB_F_ dimers *in cis* onto the DNA-tethers and these ParB_F_ dimers are capable of forming DNA looping bridges likely *via* inter-dimer interactions to form a condensed partition complex-like structure. As expected, ParB_F_^R121A^ bearing a mutation at the critical Box II residue in the CTPase domain was unable to condense DNA to a significant degree even with *parS_F_* containing tethers (Figure 6B red). We propose that stable DNA condensation by ParB_F_ is mediated by CTPase domain dimerization and requires both *parS_F_* and CTP at moderate ParB_F_ concentrations (∼100 nM) (Figure 6C).

**Figure 6.**
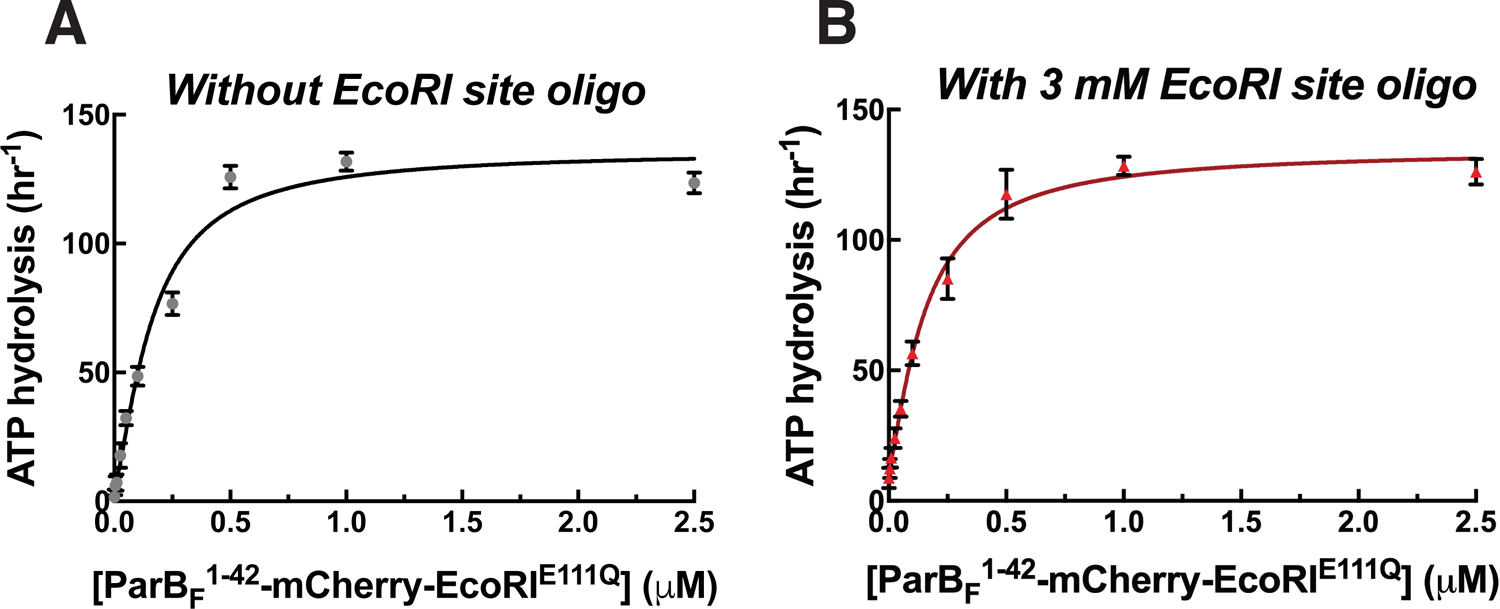
*Magnetic tweezers measurements of parS_F_ and CTP-dependent DNA condensation by ParB_F_.* (A) Schematic showing the magnetic tweezers DNA condensation assay. One end of a 5 kb DNA molecule is attached to the surface of a flow-cell and the free end is attached to a 1 µm magnetic bead (brown sphere). The DNA extension (Ex) was measured by tracking the bead height above the cover glass surface at two different forces; 0.05 pN (low force, lf), and 5 pN (high force, hf). The extent of DNA condensation was estimated from the difference in DNA extension with and without ParB_F_. (B) Changes in extension at low force (dEx^lf^ = Ex^lf^-Ex_0_^lf^), left panel, and at high force (dEx^hf^ = Ex^hf^-Ex_0_^hf^), right panel, for 7 different conditions plotted as a function of ParB_F_ concentration. The extension values were the averages of the last 5 seconds of the extension at low force (circles) and the first 5 seconds of the extension at high force (triangles). Error bars represent standard error of means (SEM). Different conditions are color coded as follows. Purple: *parS_F_*-DNA tether with WT ParB_F_ and CTP; light blue: *parS_F_*-DNA tether with WT ParB_F_ and CDP; dark blue: nsDNA tether with WT ParB_F_ and CTP; red: *parS_F_*-DNA tether with ParB_F_^R121A^ and CTP; orange: nsDNA tether with ParB_F_^R121A^ and CTP. For comparison with condensation in the presence of CTP, dEx data of *parS_F_*-DNA tether (purple) and nsDNA tether (blue) with WT ParB_F_ without CTP are displayed (inset, open circles for 0.05 pN, triangle for 5 pN respectively). (C) The condensation probabilities at 0.1 µM ParB_F_ for 5 different conditions at 0.05 pN and 5 pN. The condensation probability was calculated by dividing the number of DNA tethers that exhibited DNA condensation by the total number of DNA tethers for each measurement condition. Except for *parS_F_*-DNA with WT ParB_F_ and CTP, all conditions show either minimal or negligible condensation probabilities. The different conditions are color-coded as indicated in B and the diagonal stripes indicates probabilities at 5 pN. Error bars represent standard error of means (SEM). **Source data 1:** Numerical data for figure 6 panel A (Microsoft Excel). **Source data 2:** Numerical data for figure 6 panel B-left (Microsoft Excel). **Source data 3:** Numerical data for figure 6 panel B-right (Microsoft Excel). **Source data 4:** Numerical data for figure 6 panel C (Microsoft Excel). **Figure supplement 1—Source data 1:** Numerical data for Figure 6—figure supplement 1A (Microsoft Excel). **Figure supplement 1—Source data 2:** Numerical data for Figure 6—figure supplement 1B (Microsoft Excel). **Figure supplement 2—Source data 1:** Numerical data for Figure 6—figure supplement 2 (Microsoft Excel). **Figure supplement 3—Source data 1:** Numerical data for Figure 6—figure supplement 3 (Microsoft Excel).

**Figure 6—figure supplement 1.**
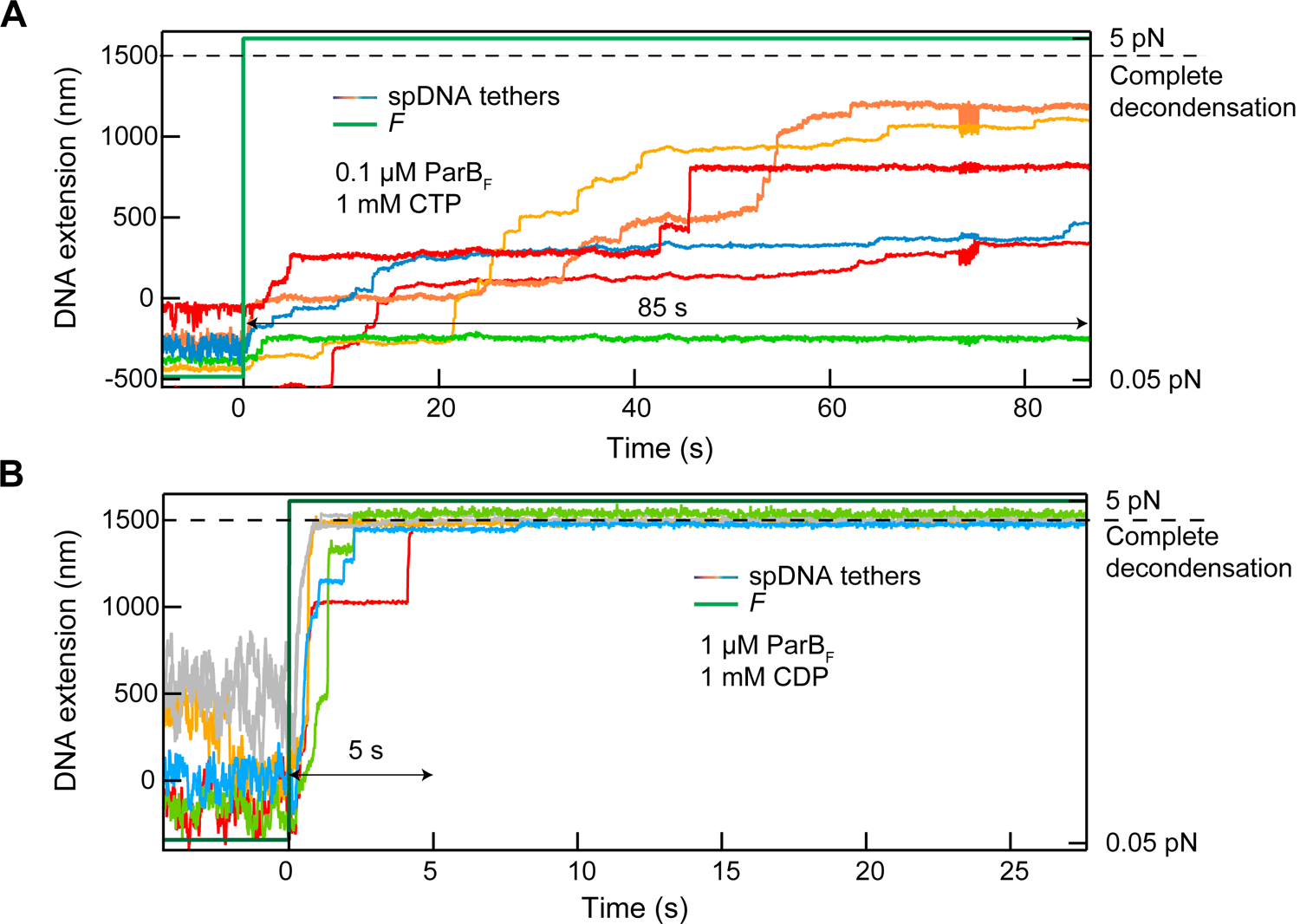
Stepwise de-condensation of condensed DNA tethers in the presence of CDP or CTP by tensile force: condensation observed in the presence of CDP is unstable. (A) The extension of 6 DNA tethers containing *parS_F_* sequences (colored solid lines, left y-axis) were simultaneously measured at a force (green solid line, right y-axis) of 0.05 pN after the introduction of ParB_F_ (0.1 µM) and CTP (1 mM). The tethers remained condensed for at least 85 s after the force was increased to 5 pN. Most increases in extension took place in discrete steps. (B) The same measurement as in (A) except that the DNA was condensed with 10-fold more ParB_F_ (1 µM) with CDP (1 mM). DNA condensation in the presence of CDP was quickly reversed under high force (5 pN). After the force was increased to 5 pN (time 0) the extension of all five DNA tethers increased to their full decondensed length (dashed line) within ∼5 s.

**Figure 6–figure supplement 2.**
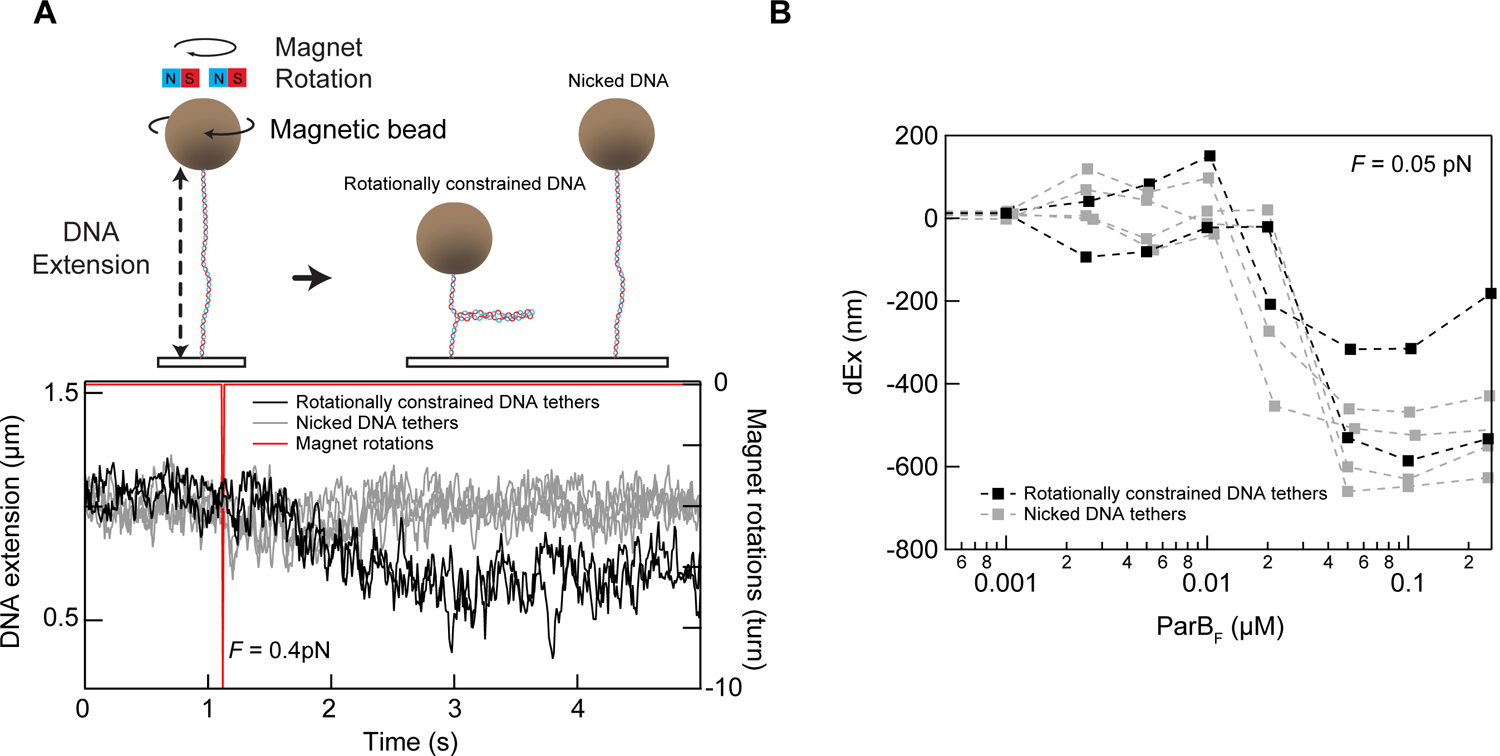
Tether condensation by ParB_F_ is comparable for topologically constrained (supercoilable) and unconstrained (nicked) DNA. (A) The DNA tethers used for the condensation experiments were a mixture of torsionally constrained and unconstrained. Prior to condensation measurements, the magnetic beads attached to the end of the DNA molecules were rotated by rotating the external magnets in the magnetic tweezers instrument. Rotationally constrained DNA molecules are supercoiled by rotating the magnet, resulting in the formation of a plectoneme and a decrease in the extension of the DNA molecule. Torsionally unconstrained (typically nicked) DNA molecules are insensitive to the bead rotation and remain at the same extension (top). An example of this *in situ* discrimination between torsionally constrained and unconstrained DNA tethers is shown in the graph displaying the extension of 6 DNA molecules (bottom). The DNA extension decreased for 2 of the 6 DNA molecules after imposing −10 turns, indicating that these two molecules are rotationally constrained (left panel bottom). (B) Condensation, as measured by the change in extension reduction as a function of ParB_F_ concentration is comparable for the topologically constrained (black squares) and unconstrained (grey squares) DNA molecules.

**Figure 6—figure supplement 3.**
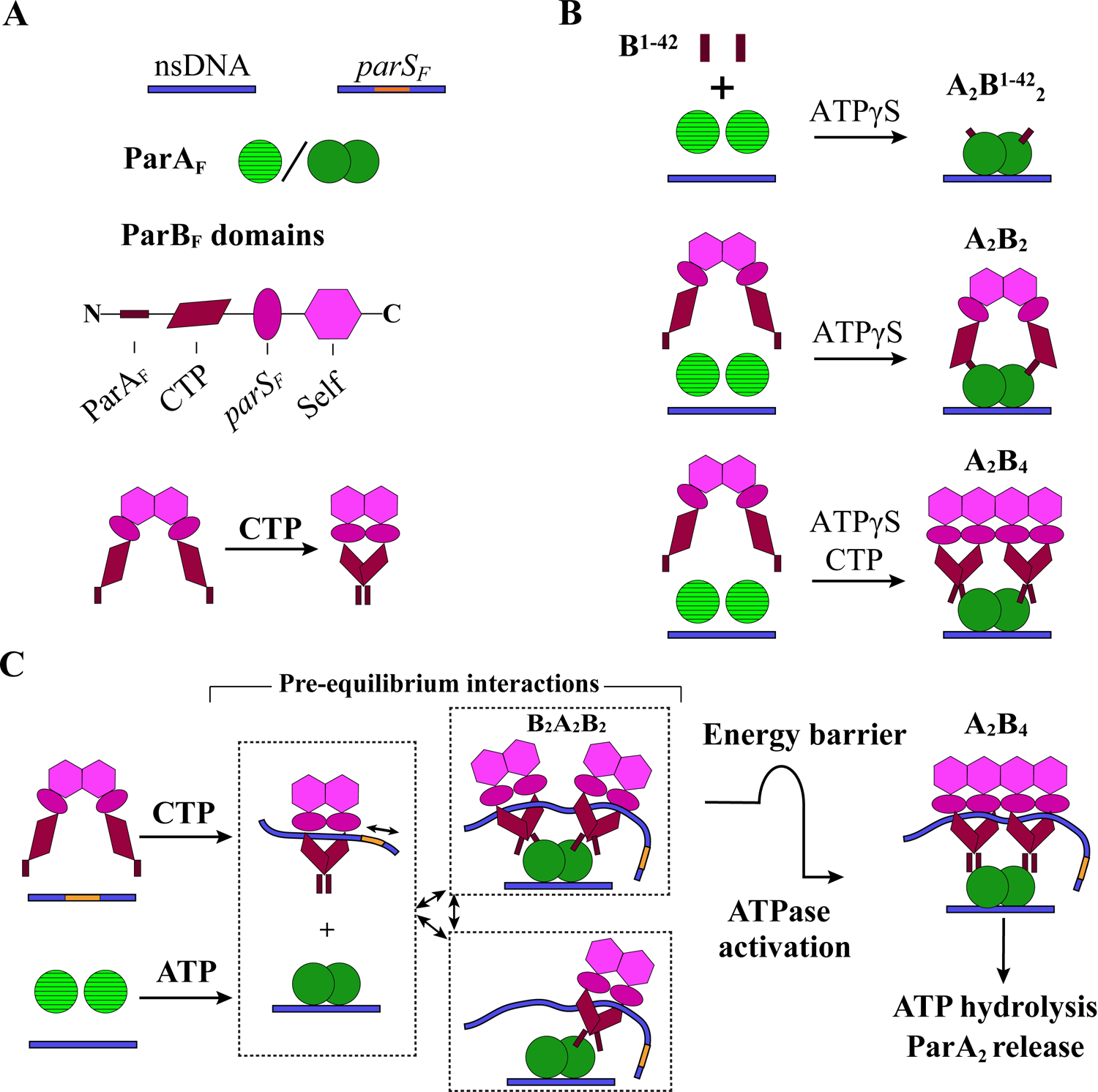
DNA tethers without *parS_F_* sequence are not condensed by ParB_F_ and *parS_F_ in trans* in the presence of CTP. DNA tethers that did not contain a *parS_F_* site were not significantly condensed by the addition of *parS_F_* DNA fragment *in trans* with ParB_F_ in the presence of CTP. The decreases in extension at 0.05 pN (blue circle, left panel) and 5 pN (blue triangle, right panel) are plotted as a function of the concentration of ParB and *parS_F_*, which were added together at stoichiometric concentrations.

## Discussion

In this report, we characterized facets of the ParA_F_—ParB_F_ interaction leading to the assembly of the nsDNA-bound ParA_F_—ParB_F_ complex that is required to activate ParA_F_ for ATP hydrolysis and dissociation from nsDNA under the influences of *parS_F_* and CTP (Summarized in Figure 7). Our results indicate that both ParB_F_ binding faces of the nsDNA-bound ParA_F_ dimers must be occupied by a ParB_F_ N-terminal domain for ATPase activation (Figure 2C-G and 7B). In principle, two copies of the ParB_F_ N-terminal domain activating a ParA_F_ dimer could belong to one ParB_F_ dimer as seen in the absence of CTP or *parS_F_* (Figure 4A-B and 7B middle). However, most ParB_F_ dimers in partition complexes *in vivo* are likely in the CTP- and *parS_F_*-activated state, spreading over a *parS_F_*-proximal DNA region. CTP or *parS_F_* binding alters the ParB_F_ dimer structure to prevent a single ParB_F_ dimer from providing both copies of the N-terminal domain to occupy both binding faces of a ParA_F_ dimer, necessitating two ParB_F_ dimers, each providing one N-terminal domain to a ParA_F_ dimer (Figure 4C-D, Figure 5 and Figure 7B bottom). Strikingly, *parS_F_* together with CTP significantly increased the A_2_B_4_ complex assembly rate on nsDNA without strongly affecting its disassembly rate. Although we have not analyzed the full kinetic details of the process that leads to ATPase activation by ParB_F_, we propose that a moderately slow transition separates formation of the ATPase-activated A_2_B_4_ complex from the rapidly reversible ParA_F_—ParB_F_ interaction processes. Such a local slow step would partially decouple the reversible ParA_F_—ParB_F_ interaction dynamics from the irreversible ATP hydrolysis, thereby promoting dynamic interactions between the nucleoid and partition complex that facilitate partitioning *via* the diffusion-ratchet mechanism as elaborated below.

**Figure 7.**
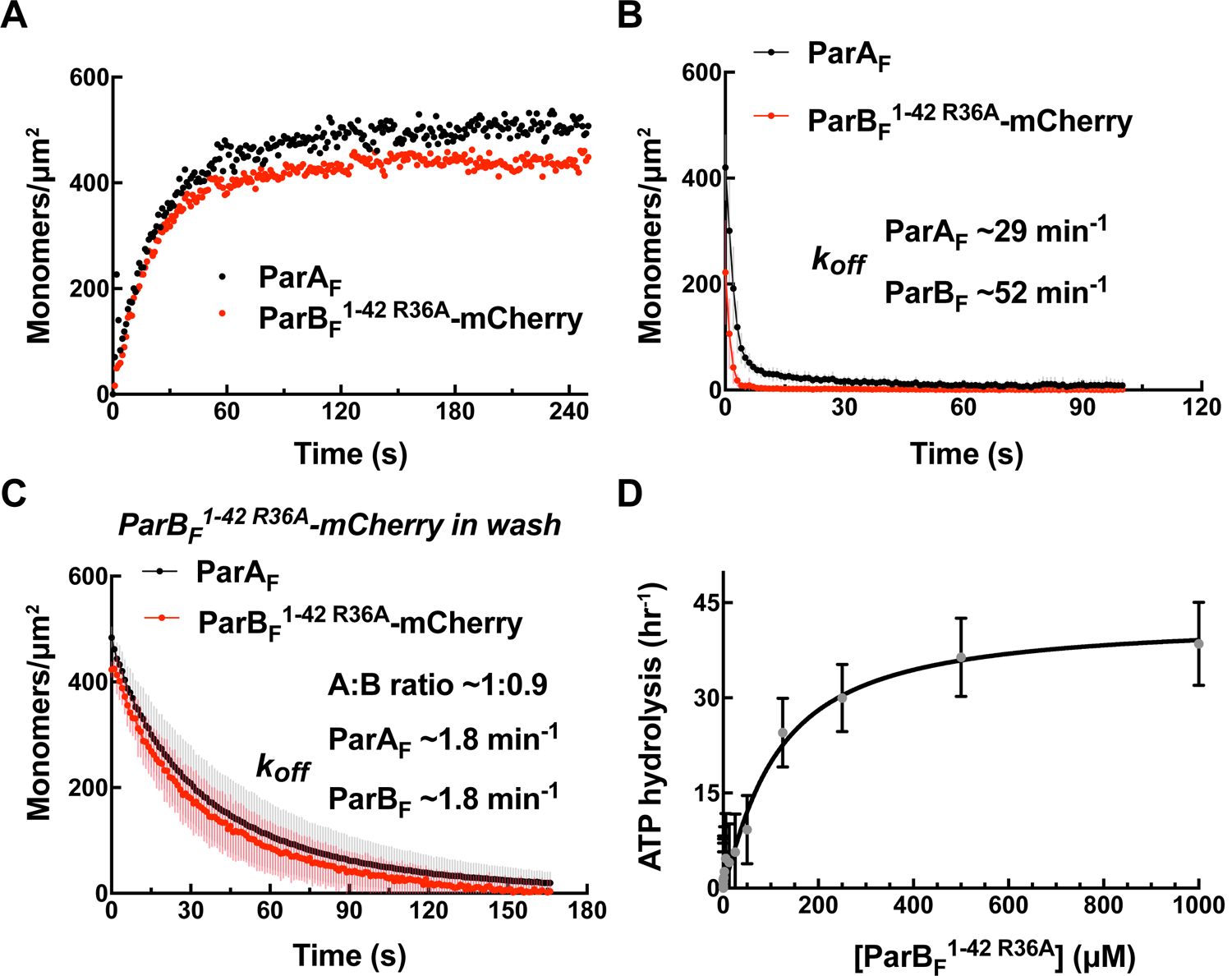
*Cartoon of the proposed pre-ATPase-activation complexes of ParA_F_ and ParB_F_*. (A) Pictograms of nsDNA, *parS_F_*-DNA, ParA_F_ monomer/dimer and ParB_F_ domains with binding ligand designations. The CTPase domains of a ParB_F_ dimer fold forming a single globular domain on binding CTP, bringing the two ParA_F_-binding domains into close proximity. (B) ParA_F_-binding domain, ParB_F_^1-42^ alone can convert ParA_F_ monomers to DNA-binding-competent dimers in the presence of ATP*γ*S by forming an A_2_B^1-42^_2_ complex (top). ParB_F_ dimers in the absence of CTP convert ParA_F_ monomers to DNA-binding-competent dimers in the presence of ATP*γ*S by straddling a ParA_F_ dimer to form an A_2_B_2_ complex (middle). In the presence of CTP, the close proximity of the ParA_F_-binding domains of the ParB_F_ dimer prevents A_2_B_2_ complex formation and instead an A_2_B_4_ complex assembles on nsDNA in the presence of ATP*γ*S (bottom). (C) In the presence of *parS_F_* and CTP, ParB_F_ dimers load on to the *parS_F_*-DNA and spread to adjacent DNA regions while adopting a state that enables faster assembly of A_2_B_4_ complexes. Considering the requirements for efficient partition complex motion by diffusion-ratchet mechanism based on the chemophoretic principle of force generation, we propose a significant energy barrier that slows the formation of the ATP hydrolysis-competent A_2_B_4_ complex. This energy barrier partially decouples ParA_F_—ParB_F_ association-dissociation dynamics from ATP hydrolysis, which triggers ParA_F_ dissociation from the nucleoid.

The clearest indication that both ParB_F_-interacting faces of the nsDNA-bound ParA_F_ dimer must be occupied by the N-terminal domain of ParB_F_ for ATPase activation came from experiments using artificial ParB_F_ constructs. We showed that monomeric ParB_F_^1-42^ stimulates ParA_F_ ATPase with a clear sigmoidal concentration dependence, indicating that one molecule of ParB_F_^1-42^ binding to one side of a ParA_F_ dimer cannot fully activate the ParA_F_ ATPase (Figure 2F). When ATP hydrolysis was blocked by using non-hydrolysable ATPγS, ParB_F_^1-42^-mCherry formed an equimolar complex with ParA_F_ (Figure 2C-E). Thus, the ParA_F_ forms a complex with ParB_F_^1-42^-mCherry bound at both ParB_F_-interacting faces of the ParA_F_ dimer prior to ATP hydrolysis (Fig 7B; top). Consistently, an artificially dimeric ParB_F_^1-42^ construct, ParB_F_^1-42^-mCherry-EcoRI^E111Q^, efficiently activated ParA_F_-ATPase with hyperbolic concentration dependence (Figure 2G).

In the absence of CTP or *parS_F_*, the ParB_F_ dimer is held together by the C-terminal self-dimerization domain (Fig 7A bottom) with dimerization *K_D_* of ∼19 nM (Figure 4—figure supplement 1). In this state, the N-terminal halves of the monomers are thought to be separate from each other according to the SAXS envelope of the structure (Chen et al., 2015; also see Figure 2— figure supplement 3B). Thus, one dimer could straddle a ParA_F_-dimer with each N-terminal ParA_F_-interaction domain interacting with one of the two faces of a ParA_F_-dimer (Figure 7B middle).

In contrast, when bound by CTP the CTPase domains in a ParB_F_ dimer fold to form a single globular domain (Figure 7A bottom) (Soh et al., 2019; see Figure 2—figure supplement 3C). The two ParA_F_-interaction domains emanating from this dimeric domain are unlikely to reach both sides of a ParA_F_ dimer, necessitating an A_2_B_4_ complex for ATPase activation (Figure 7B bottom). In theory, it is possible that a chain of (A_2_B_2_)_n_ might form, but the 1:2 protein stoichiometry observed in the presence of ATP*γ*S indicates such a configuration is unfavored. A_2_B_4_ complexes formed in the presence of CTP and *parS_F_* DNA fragments contained almost no *parS_F_* DNA fragments (Figure 5D). This is consistent with the notion that after binding *parS_F_*, CTP-ParB_F_ dimers convert to a low *parS_F_*-affinity state while remaining topologically bound to the DNA and spreading to surrounding DNA regions (Soh et al., 2019). The A_2_B_4_ complex formed in the presence of *parS_F_* DNA fragments, even without CTP, contained significantly less than a stoichiometric amount of *parS_F_* fragment (Figure 5D). This suggests that association with nsDNA-bound ParA_F_ dimer lowers the affinity of ParB_F_ for *parS_F_*, perhaps shifting the structure closer toward *parS_F_*-activated ParB_F_-CTP.

The ParB: ParA stoichiometry change from 1:1 to 2:1 caused by *parS_F_* (Figure 4C, D) did not occur with the Box II mutant ParB_F_^R121A^ (Figure 4—figure supplement 2). It is possible that when a ParB_F_ dimer binds *parS_F_*, the adjacent CTPase domains of the two monomers adopt a mutually interacting folded state akin to the CTP-bound state even without CTP, promoting a BoxII-dependent dimerized domain structure. This may disfavor formation of the A_2_B_2_ complex, favoring the A_2_B_4_ complex that was observed.

All the A_2_B_2_ and A_2_B_4_ complexes we observed in the presence of ATP*γ*S dissociated from nsDNA more slowly (k_off_ = 0.5 - 1 min^-1^; Supplementary Table 1) compared to ParA_F_ in the absence of ParB_F_ (∼6 min^-1^; Fig 2B). For the case of monomeric ParB_F_^1-42^, which dissociated from ParA_F_ more rapidly, the presence of 10 μM ParB_F_^1-42^ in the wash buffer restored the low apparent nsDNA dissociation rate constant of the complex (Figure 2E). Nevertheless, ParA_F_ dimers in these complexes appear to be primed for further conformational change toward less stably DNA-associated state. Upon dissociation of ParB_F_^1-42^ from the A_2_B^1-42^_2_ complex, ParA_F_ dissociated from the DNA-carpet within a second or so, much faster than the ATP*γ*S-ParA_F_-dimer that has not yet formed a A_2_B_2_ or A_2_B_4_ complex (Figure 2D, 3B).

Our single-molecule DNA condensation measurements indicate that CTP-bound ParB_F_ dimers are activated by contacting *parS_F_* to load *in cis* onto the *parS_F_*-carrying DNA in numbers exceeding the copy number of the *parS_F_*-consensus sequence (ParB spreading) as shown by others for chromosomal ParBs (Jalal et al., 2020; Soh et al., 2019), and condense the DNA forming an *in vivo* partition complex-like structure (Figure 6). Although the magnetic tweezers instrument used in this study did not allow direct measurement of the number of ParB_F_ molecules contained in the condensed DNA, the large number of de-condensation steps observed when high tension was applied is consistent with the presence of a large number of ParB_F_ dimers in the condensed DNA (Figure 6, Figure 6—figure supplement 1A). CDP failed to support efficient condensation of *parS_F_*-carrying DNA by ParB_F_ and the limited condensation observed, which could be due to the contaminating material in the CDP used, was disrupted far more readily than CTP-supported condensates (Figure 6, Figure 6—figure supplement 1B). Our results show that DNA-condensation is caused by ParB_F_—ParB_F_ interactions forming DNA-looping bridges without requiring other protein factors. Combined with evidence indicating that *parS_F_*-activated ParB_F_—CTP adopts a unique conformational state (Soh et al., 2019), we favor the view that DNA-bridging capability, mediated by inter-dimer ParB_F_ interaction, is another attribute of this ParB_F_ state.

Our observation indicates that the state of ParB_F_ discussed above is maintained after release from *parS_F_*-containing DNA. ParB_F_ associates with nsDNA-bound ParA_F_ dimers forming the A_2_B_4_ complex with a faster apparent assembly rate in the presence of *parS_F_* and CTP together than with either CTP or *parS_F_* alone. This observation is consistent with the decreased half-saturation concentration in the ATPase activation assay (Figure 5F, Table 2). According to the sliding clamp model of spreading ParB—CTP dimers proposed by Soh et al., (2019), ParB_F_—CTP dimers loaded onto a short *parS_F_* DNA fragments would quickly slide off the DNA as shown by Jalal et al., (2020). Since our ATPase activation assay and the DNA-carpet-bound A_2_B_4_ complex assay in the presence of CTP and *parS_F_* were done using a short linear *parS_F_* DNA fragment, the *parS_F_*-activated state of the ParB_F_—CTP dimers we described in this study must remain in this “activated” state for an extended period after sliding off the *parS_F_* fragment. Accordingly, the A_2_B_4_ complexes bound to the DNA-carpet in the presence of CTP, ATP*γ*S and *parS_F_* fragments contained almost no *parS_F_* fragments (Figure 5D). Thus, *parS_F_* acts as a catalyst to convert ParB_F_—CTP dimers from a pre-activation state to an activated state capable of faster A_2_B_4_ complex assembly. This notion is also consistent with the observation that significantly less than a stoichiometric concentration of *parS* DNA relative to ParB is sufficient for full activation of the ParB CTPase (Figure 5—figure supplement 2C; Soh et al., 2019). Although the ParB_F_ dimers in this activated state failed to load efficiently onto DNA lacking *parS_F_* sequences *in trans* (Figure 6—figure supplement 3), in the absence of contrary evidence, the parsimonious assumption is that this ParB_F_ dimer retains the conformation of spreading ParB_F_ dimers that remain loaded on the *parS_F_*-containing DNA *in cis*. Thus, we propose that the functional properties of ParB_F_ we observed in the presence of CTP and *parS_F_*, both in facilitating assembly of A_2_B_4_ complexes and activating ParA_F_-ATPase, reflect those of the majority of ParB_F_ dimers in partition complexes *in vivo*.

Our study, together with previous studies, indicates that the ATP turnover rates of ParABS systems are slow because of multiple, slow kinetic steps. These slow steps are strategically placed in the reaction pathway in order to tune the system and drive the motion of the partition complex through the diffusion-ratchet mechanism (Sugawara and Kaneko, 2011; Vecchiarelli et al., 2010). Even at saturating concentrations of ParB_F_ in the *parS_F_*-activated CTP-bound state, the maximum ATP turnover rate of ParA_F_ remained modest (∼80 ATP/ParA_F_-monomer/hour; Figure 5). The slow reactivation of ParA nucleoid binding after ATP hydrolysis likely dominates the overall ATPase cycle time (Vecchiarelli et al., 2010). The presence of a large fraction of ParA_F_ in DNA-unbound state during the steady state ATPase assay was evidenced by the fact that the half-saturation concentration of ParB_F_ (in the presence of CTP and *parS_F_*) forming the A_2_B_4_ complex was ∼0.2 μM while the total ParA_F_ concentration was 1 μM, suggesting less than ∼20% of ParA_F_ was in the nsDNA-bound state ready to interact with ParB_F_. *In vivo* the reactivation rate is likely lower since nucleoid-bound ParA is only fully activated on encountering the partition complex, which lowers the concentration of ParA in the cytosol waiting to be reactivated. The lower precursor concentration slows the nucleoid rebinding rate of ParA non-linearly because reactivation involves a relatively fast nucleotide-dependent reversible ParA dimerization with apparent *K_D_* of ∼2 μM, followed by a slow conformational step. This makes the process dimerization-limited at lower precursor ParA concentrations according to the study of ParA_P1_ (Vecchiarelli, 2010). Whereas this slow ParA reactivation and rebinding process, which allows the maintenance of the nucleoid-bound ParA concentration gradient (Hu et al., 2017), is a critical element of chemophoresis driven motility, the rate of ParA-ATPase activation by ParB is another important factor. In particular, efficient chemophoresis force generation relies on ParA_F_—ParB_F_ interactions achieving a local quasi-equilibrium prior to ATP hydrolysis (Sugawara and Kaneko, 2011). Therefore, we speculate that there is a significant energy barrier associated with the conformational transition of a ParA_F_— ParB_F_ complex to achieve ATPase activation (Figure 7C). The resulting local time delay, in addition to the fact that two ParB dimers are required to bind a ParA dimer to activate its ATPase, would partially decouple the pre-ATP hydrolysis ParA_F_—ParB_F_ reversible interaction steps from the ATP hydrolysis step. This delay would in turn permit ParB—ParA binding to approach local quasi-equilibrium, increasing the efficiency of ParA distribution gradient sensing and motive force generation by the partition complex. In addition, this slow activation step would prevent possible over-depletion of the local nucleoid-bound ParA_F_ as the partition complex establishes the ParA_F_ depletion zone.

Disassembly of the ATP-bound A_2_B_4_ complex might be slow prior to ATP hydrolysis considering the stability of the complexes in the presence of ATP*γ*S. Thus, we propose the energy barrier postulated above is positioned immediately prior to formation of this complex rather than between this complex assembly and ATP hydrolysis. A slow step after formation of the stable complex would prolong the lifetime of the link between the nucleoid and the partition complex impeding partition complex motion without permitting the reversible ParA_F_—ParB_F_ interaction to approach equilibrium. We consider this conformational transition is likely the step synergistically accelerated by CTP and *parS_F_*. We note that CTP-activated ParB_F_ stimulates ParA_F_ ATPase with sigmoidal concentration dependence (Figure 5E-F, Table 2) suggesting two ParB_F_ dimers separately bind a ParA_F_ dimer during a pre-equilibrium binding phase, forming a transient B_2_A_2_B_2_ complex. We imagine the slow conformational step proposed here might be assisted by the property of the CTP/*parS_F_*-activated ParB_F_ dimers that promotes inter-dimer interactions as suggested by the magnetic tweezers experiments, stabilizing the interaction between the two ParB_F_ dimers within a complex, depicted as conversion of B_2_A_2_B_2_ complex to A_2_B_4_ complex in Figure 7C. This might explain the higher assembly rate and stability of the complex formed with *parS_F_*-activated ParB_F_-CTP. Yet, CTP and *parS_F_* DNA do not significantly increase the ATP turnover rate of ∼80 h^-1^, indicating that the proposed kinetic delay time must be a small fraction of the ATPase cycle time (∼45 sec), for which we believe the rate limiting step resides in the reactivation process of ParA for nsDNA binding after ATP hydrolysis (Vecchiarelli et al., 2010). Assembly of the A_2_B^1-42^_2_ complex perhaps does not experience this time delay due to fewer steric constraints, but ParB_F_^1-42^ dissociates more readily compared to full-length ParB_F_. If two CTP-bound and parS_F_-activated ParB_F_ dimers independently associating with a nucleoid-bound ParA_F_-ATP dimer is important for efficient partition complex motive force generation by the chemophoretic principle as proposed above, one might be able to design a mutant ParB_F_ that can activate ParA_F_-ATPase by forming an A_2_B_2_ complex even in the presence of CTP, which would significantly affect plasmid partition efficiency. Efforts to generate such ParB_F_ mutants are currently under way.

This study demonstrates how *parS_F_*, along with CTP, has wide-reaching roles in the F-plasmid ParABS system; not only in ParB_F_’s ability to spread from *parS_F_* and promote ParB_F_—ParB_F_ interactions for partition complex compaction, but also in ParB_F_ dimer interactions with ParA_F_. However, we still need to investigate how the ParB_F_ CTPase activity is impacted by *parS_F_* in different states of the ParB_F_—*parS_F_* complex and its interaction with the ParA_F_-DNA complex. More generally, in order to understand how the system is orchestrated to achieve system dynamics that result in robust plasmid segregation, improved understanding of the microscopic kinetic parameters is essential. Many details of the system dynamics still remain to be addressed to understand the full picture of the ParABS partition mechanism.

## Materials and Methods

### Key Resources Table

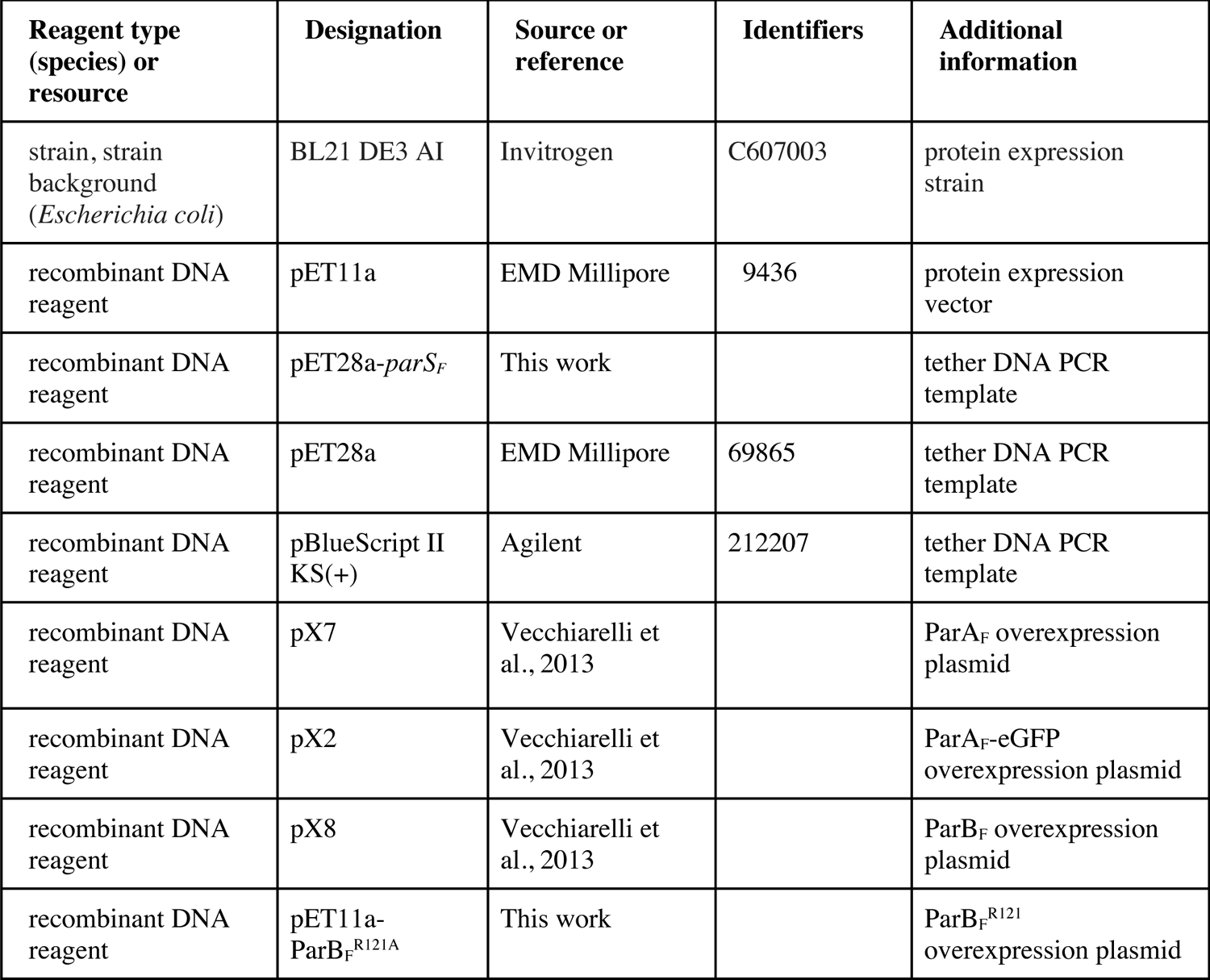
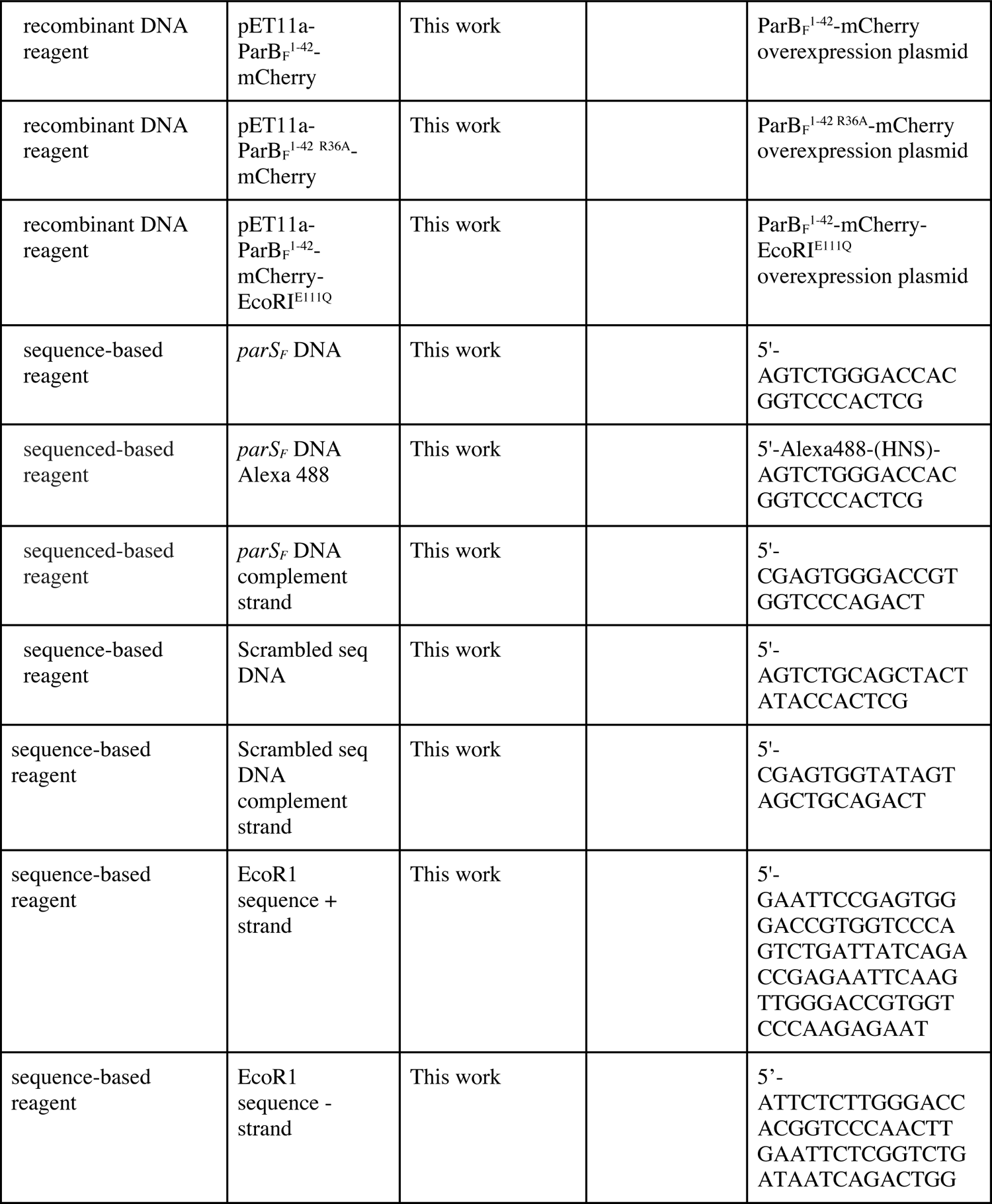
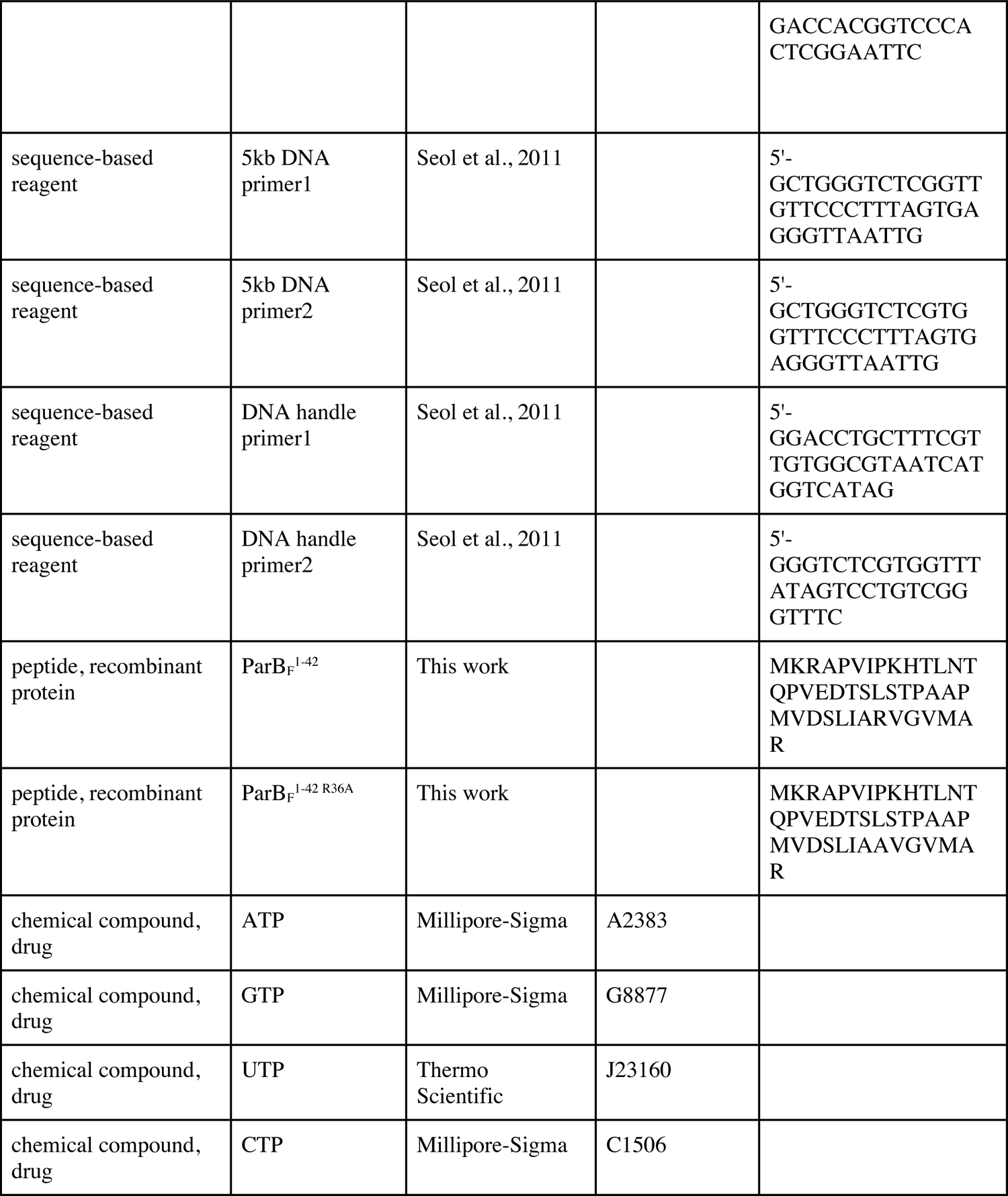
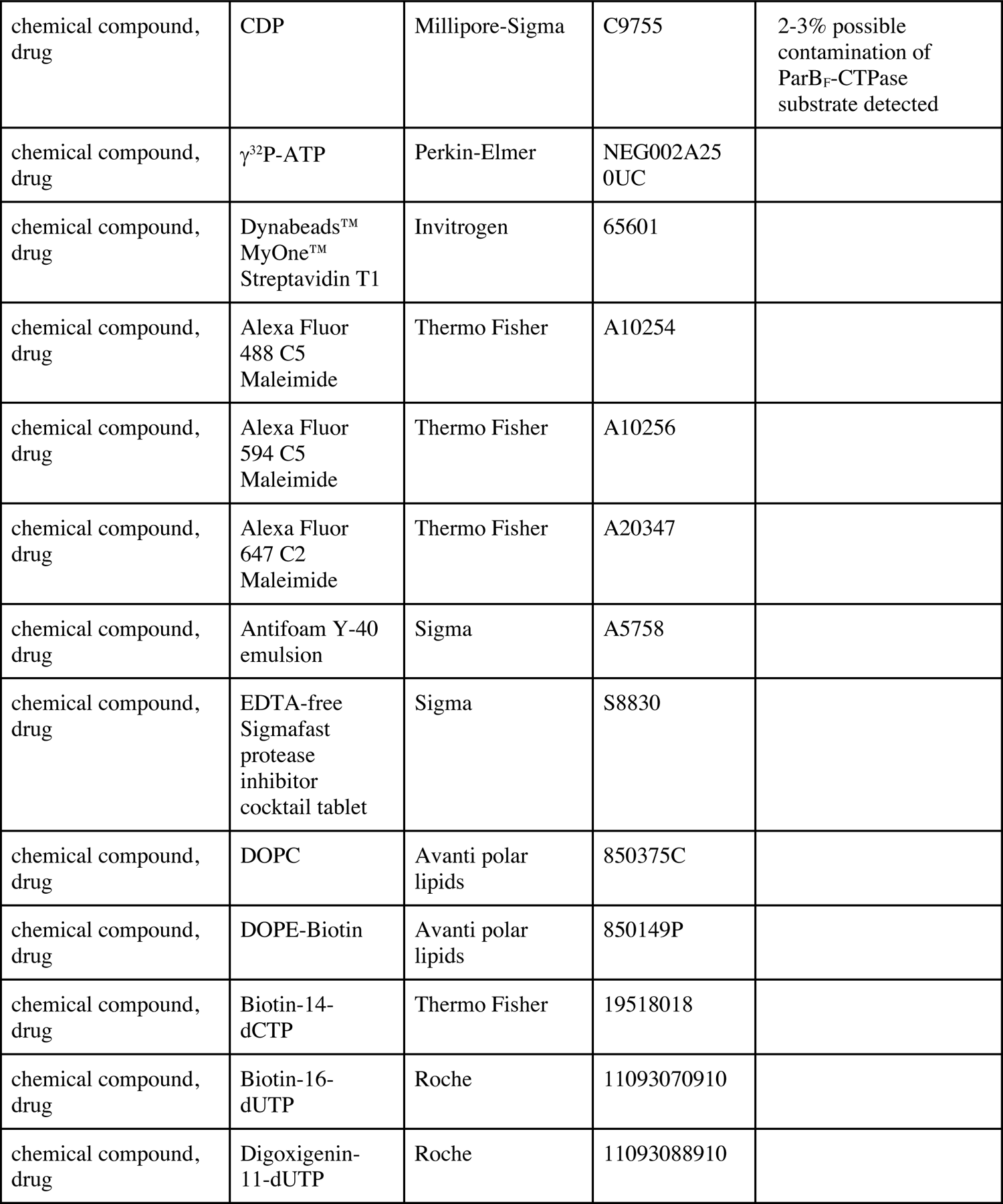
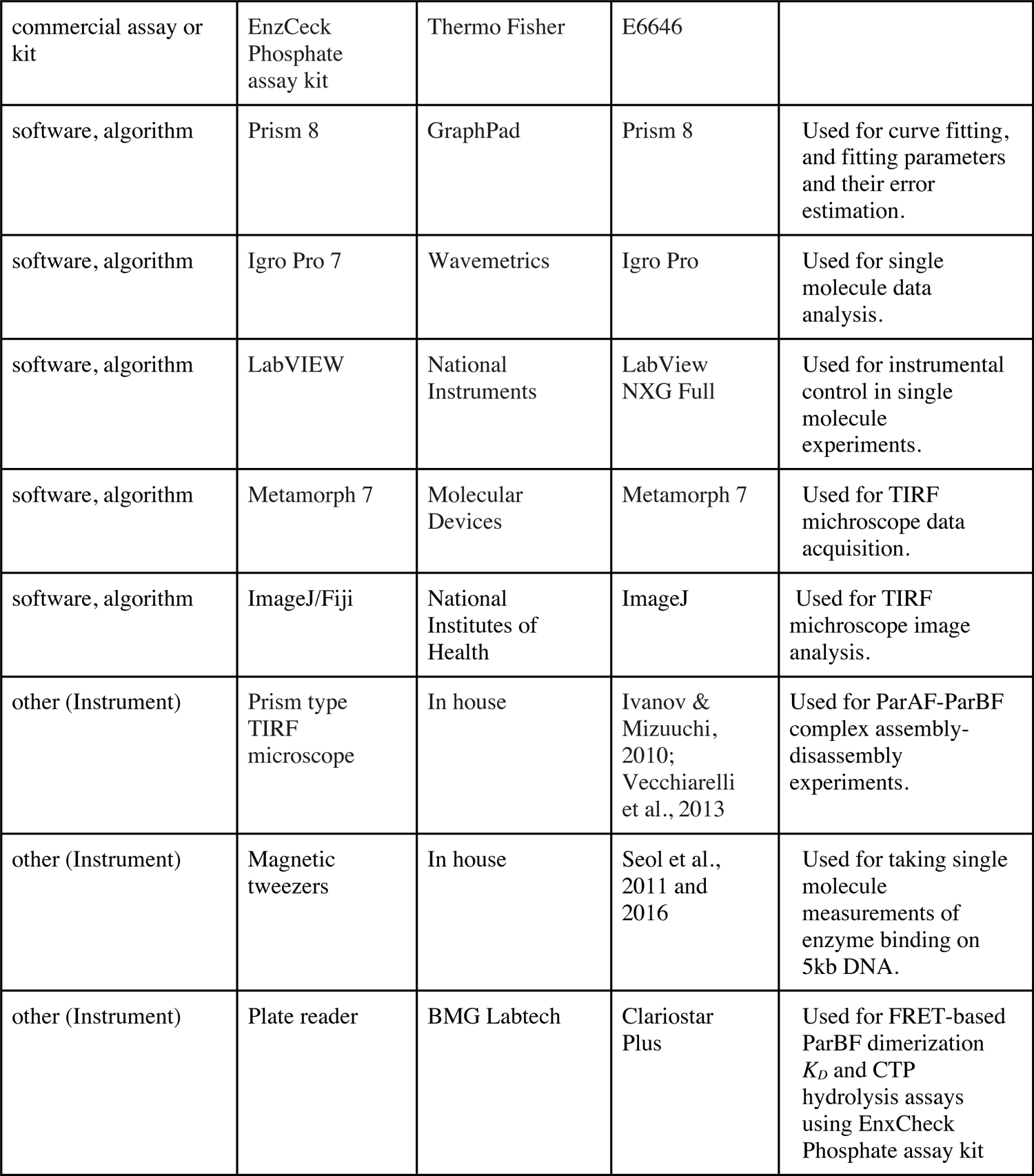

### Plasmids and constructs for protein expression

All expression open reading frames were synthesized and subcloned into pET11a (Genscript). ParA_F_, ParA_F_-eGFP, ParB_F_, ParB_F_^R121^, ParB_F_^1-42^-mCherry, ParB_F_^1-42 R36A^-mCherry and ParB_F_^1-42^-mCherry-EcoRI^E111Q^ constructs were made with a hexa-histidine tag on their C-terminus. Protein fusions were made with a SGGG linker between fused domains, with exception of ParB_F_^1-42^-mCherry-EcoRI^E111Q^ which had a 4x (SGGG) linker between ParB_F_^1-42^ and mCherry. ParB_F_^1-42^ and ParB_F_^1-42 R36A^ were synthesized *de novo* (Genscript).

### Oligonucleotides

The 24 bp double-stranded DNA fragments containing the *parS_F_* consensus sequence and a scrambled sequence used in this study were: 5’-AGT CTG GGA CCA CGG TCC CAC TCG; 5’ - AGT CTG CAG CTA CTA TAC CAC TCG, respectively and their complements. The fluorescently labelled *parS_F_* substrate was synthesized with Alexa-488 NHS coupled to the 5’ of the forward strand by the manufacturer (IDT).

### Protein purification and fluorescent labelling

For expression of proteins 5 ml of an overnight culture of BL21 DE3 AI (Invitrogen) *E. coli* cells transformed with the desired plasmid were inoculated into 500 ml Terrific Broth (Teknova) supplemented with 100 μg/ml carbenicillin, antifoam Y-40 emulsion (Sigma), 1 g/l NaCl, 0.7 g/l Na_2_SO_4,_ 2.6 g/l NH_4_Cl, and 0.24 g/l MgSO_4_. The cultures were incubated at 37 °C in 2.5 l Fernbach flasks and shaken at 120 rpm until they reached an OD_600_ of 1.8. Cultures were chilled on ice before they were induced by the addition of 1 mM IPTG and 0.2 % L-arabinose. Following induction, cultures were incubated at 16 °C for 16 hr and cells were harvested by centrifugation at 6000 x g for 15 min at 4 °C. Cell pellets were frozen in liquid nitrogen and stored at −80 °C.

Frozen cells were thawed and resuspended to a density of 1 g cell pellet/10 ml in lysis buffer (ParA_F_, ParA_F_-eGFP and ParB_F_^1-42^-mCherry-EcoRI^E111Q^: 25 mM Tris.HCl pH 8, 1 M NaCl, 20 mM imidazole, 2 mM β-mercaptoethanol, 10% glycerol; ParB_F_ and ParB_F_^R121A^: 10 mM Sodium Phosphate buffer pH 7, 1 M NaCl, 20 mM Imidazole, 2 mM β-mercaptoethanol, 10% glycerol; ParB_F_^1-42^-mCherry and ParB_F_^1-42/R36A^-mCherry: 25 mM HEPES.KOH pH 7.5, 150 mM NaCl, 6 M guanidinium chloride, 20 mM Imidazole, 2 mM β-mercaptoethanol, 10% glycerol) containing EDTA-free Sigmafast protease inhibitor cocktail tablet (Sigma) using a homogenizer. Lysozyme and Benzonase (Sigma) were added to a concentration of 1 mg/ml and 50 u/ml, respectively, and the cells were lysed *via* a microfluidizer. Cell debris were pelleted by centrifugation at 142,000 x g for 45 min at 4 °C, and the supernatant passed through a 0.22 μm filter.

Lysate was loaded on to a 5 ml HisTrap HP cassette (GE Healthcare) equilibrated in lysis buffer. The cassette was then washed with 10 column volumes of lysis buffer followed by 10 column volumes HisTrap buffer (as lysis buffer without guanidinium hydrochloride and with the following NaCl concentrations: ParA_F_ proteins, 200 mM; ParB_F_ proteins, 150 mM), and the protein eluted with a gradient from 20 to 500 mM imidazole over 10 column volumes using an AKTA Pure (GE Healthcare).

All proteins except ParB_F_^1-42^-mCherry and ParB_F_^1-42 R36A^-mCherry were then subjected to ion-exchange chromatography. The peak fractions from the HisTrap column were pooled and slowly diluted whilst stirring with a Mono Q/S-buffer (as lysis buffer without imidazole or NaCl, but with 0.1 mM EDTA pH 8) until the conductivity of the sample was: 18 mS/cm for ParA_F_ proteins, 5 mS/cm for ParB_F_^1-42^-mCherry-EcoRI^E111Q^ and 15 mS/cm for all other ParB_F_ proteins. The conductivity of the samples was monitored using a conductivity meter (Hanna). The sample was loaded onto either a 1-ml Mono Q (ParA_F_ proteins and ParB_F_^1-42^-mCherry-EcoRI^E111Q^) or Mono S (other ParB_F_ proteins) 5/50 GL ion exchange column (GE Healthcare) pre-equilibrated with Mono Q/S-buffer containing a NaCl concentration to match the conductivity of the sample. The column was then washed with 10 column volumes of Mono Q/S-buffer + NaCl. The protein was eluted with a gradient up to 500 mM NaCl over 10 column volumes.

Finally, all protein samples were purified by size-exclusion chromatography. The peak fractions from the previous column were pooled and diluted 50:50 with concentration buffer (25 mM HEPES.KOH pH 7.5, 2 M NaCl, 2 mM β-mercaptoethanol, 10% glycerol) and concentrated to ∼2 ml using a Centriprep 10-kDa spin concentrator (Millipore). The sample was then injected onto an S200 16/600 size exclusion column (GE Healthcare) pre-equilibrated in gel filtration buffer (25 mM HEPES-KOH pH 7.5, 0.1 mM EDTA pH 8, 0.5 mM TCEP and 10% glycerol with 600 mM KCl for ParA_F_ proteins and 150 mM KCl for ParB_F_ proteins). Peak fractions were then pooled and concentrated to ∼100 µM (∼5-10 mg/ml) as determined by UV 280 nm absorbance before being aliquoted, frozen in liquid nitrogen and stored at –80 °C. Protein aliquots were used once and not subjected to freeze-thaw cycles.

To produce fluorescently labeled ParB_F_ and ParB_F_ ^R121A^, ParB_F_ protein was buffer exchanged into gel filtration buffer without reducing agent and incubated with a two-fold molar excess of Alexa Fluor 647 C2 Maleimide (Thermo Fisher) for 30 min at room temperature. The reaction was then quenched by the addition of DTT to a final concentration of 10 mM. The protein solution was then filtered through a 0.22 μm filter and free dye removed by buffer exchange into gel filtration buffer in Amicon ultra 10 kDa spin concentrator (Millipore). The extent of labeling was estimated based on absorbance at 280 and 647 nm.

### Assaying contaminating activities in the protein preparations

Proteins purified by the above protocol had no significant DNA endonuclease activity. After 16 hr incubation of supercoiled pBR322 with 2 µM ParA_F_ and/or 10 µM ParB_F_ at 37 °C in ATPase buffer (see below) no linear DNA was observed and less than 10% of the supercoiled plasmid was converted to a nicked-circular form. The contaminating ATPase activity for all ParB_F_ proteins was less than two mol ATP per mol ParB_F_ per hour, as determined by the ATPase assay protocol detailed below.

### ATPase activity assays

Steady-state ATPase activity was measured as described (Vecchiarelli et al., 2016) with modifications. ATP to be used for ATPase activity assays was purified after diluting 20 μCi ATP γ-P^32^ (Perkin-Elmer) in 100 µl of 100 mM unlabeled ATP (Sigma) by passing through a 3 ml P2 resin size-exclusion column equilibrated with a buffer containing 50 mM HEPES·KOH pH 7.5, 150 mM KCl and 0.1 mM EDTA. The purity of fractions was determined by TLC. 1 μl of each fraction was spotted on to a 10 x 8 cm piece of TLC PEI Cellulose F paper (Millipore) 1 cm above the bottom of the paper and developed for 10 mins using 400 mM NaH_2_PO_4_ pH 3.6 as the solvent. The fractions containing the minimum contamination of P^32^-Pi were pooled and their concentration determined by spectrometry before storage at −20 °C.

ParA_F_ ATPase activity was measured in the presence of the combinations and concentrations of proteins and DNA cofactors specified in the main text in ATPase buffer (50 mM HEPES·KOH pH 7.5, 150 mM KCl, 5 mM MgCl_2_, 0.5 mM TCEP and 1 mM ATP γ-P^32^). Reactions were incubated at 37 °C for 4 hrs and stopped by the addition of an equal volume of 1 M formic acid. The increase of P^32^-Pi was measured by TLC using PEI Cellulose F paper as detailed above.

### ParB_F_ NTPase activity assays

Steady state ParB_F_ CTPase activity was measured in CTPase buffer containing 50 mM Tris-HCl pH 7.5, 100 mM NaCl, 2 mM MgCl_2_, 1 mM DTT, 100 μg/ml BSA, 200 μM MESG (EnzChek probe), 1 U/ml of purine nucleotide phosphorylase, and ParB_F_, *parS_F_* DNA and CTP at concentrations specified in the figure, following the protocol of the supplier of the EnzChek phosphate assay kit (ThermoFisher). Reactions were typically repeated three times using 96-well microtiter plates and the 360 nm absorption signal increase was monitored at half to one min intervals using Clariostar Plus plate reader (BMG Labtech). The absorption signal increase after subtraction of background time course in the absence of enzyme was converted to released Pi concentration increase based on phosphate titration measurements. The CTP hydrolysis rate was calculated from the initial slope of the time course curve, which typically started after ∼7 min deadtime for the plate setting up. Substrate specificity was examined comparing Pi release from four ribonucleoside triphosphates. Attempt to examine inhibition of the CTPase activity by CDP or to detect CDP binding to ParB_F_ was postponed when the CDP used in this study was found to release Pi upon incubation with ParB_F_. CDP obtained from two additional suppliers also generated similar quantities of Pi upon incubation with ParB_F_.

### TIRF microscopy

The general design of the TIRF microscopy setup was essentially as previously described (Ivanov and Mizuuchi, 2010; Vecchiarelli et al., 2013). A prism-type TIRFM system was built around an Eclipse Ti microscope (Nikon) with a 40x objective (S Fluor, 40x/1.30 oil, Nikon) and two-color images captured by an Andor DU-897E camera through a dxcr630 insert DualView (Photometrics) with the following settings: 3 MHz digitizer (grey-scale); 5.2 pre-amplifier gain, 2 MHz vertical shift speed; +1 vertical clock range; electron-multiplying gain 30; EM CCD temperature set at −90 °C; baseline clamp ON; and exposure time 100 ms.

The excitation for ParA_F_-eGFP and Alexa647-ParB_F_ were provided by a 488 nm diode-pumped solid-state laser (Sapphire, Coherent) and a 633 nM HeNe laser (Research Electro-Optics), respectively. The TIRF illumination had an elliptical Gaussian shape in the field of view therefore intensity data for DNA-carpet-bound ParA_F_-eGFP and Alexa647-ParB_F_ signals were taken at or near the middle of the illumination profile.

Movies were acquired using Metamorph 7 (Molecular Devices) and transferred to ImageJ (National Institutes of Health) for analysis.

Flow cells were assembled using fused silica microscope slides with pre-drilled inlet/outlet ports (Esco products), #1 glass cover slips (24 x 50 mm, Thermo Fisher) and 0.001”-thick acrylic transfer tape (3M). The fused silica slide was cleaned by soaking overnight in a solution of Nochromix (Sigma)-sulfuric acid, followed by extensive rinsing with de-ionized water, drying by blowing nitrogen gas, followed by oxygen plasma treatment (South Bay Technology Inc). The Y-shaped flow path pattern was cut out of the transfer tape using a laser cutter before the flow cell assembly. Nanoports (Idex) were attached to the fused silica slide for the inlet and outlet tube connections using Norland Optical Adhesive (Thorlabs), cured by 365 nm UV light. The assembled flow cells were then baked at 80 °C with gentle compression for 2 hr.

To assemble a DNA-carpet in a flow cell, small unilamellar vesicles (SUVs) of 1,2-dioleoyl-*sn*-glycero-3-phosphocholine (DOPC) and 1,2-dioleoyl-*sn*-glycero-3-phosphoethanolamine-N-(biotinyl) (DOPE-Biotin) (Avanti Polar Lipids) were prepared as follows. 0.5 ml of DOPC (25 mg/ml chloroform) was mixed with 5 μl of DOPE-biotin (25 mg/ml chloroform) in a glass test tube and most of the solvent removed *via* evaporation under a nitrogen flow. The remaining solvent was removed by drying in a SpeedVac (Savant) at 42 °C for 1 hr followed by a further 1 hr at room temperature. 2.5 ml of degassed TK150 buffer (25 mM Tris.HCl pH 7.5, 150 mM KCl) was then added and the lipids stored, covered under nitrogen gas overnight. The lipids were then resuspended by vortexing and sonicated (70-80 watts, 30 sec on, 10 sec off) in a cup horn with water chiller set to 16 °C (QSonica) until transparent. The resulting solution of SUVs was then filtered through a 0.22 μm filter, aliquoted and stored under nitrogen gas at 4 °C for up to 4 weeks.

To prepare biotinylated salmon sperm DNA for DNA-carpets 10 mg/ml salmon sperm DNA (Thermo Fisher) was sonicated for 5 min (110 watts, 10 sec on, 10 sec off) to produce short fragments. Sonicated salmon sperm DNA was then diluted to 1 mg/ml in Terminal Transferase buffer (NEB) with 0.25 mM CoCl_2_, 40 μM Biotin-14-dCTP (Thermo Fisher) and 1 unit/μl Terminal Transferase (NEB). The DNA was incubated at 37 °C for 30 min, then the reaction stopped by heat inactivation at 75 °C for 20 min. Free Biotin-14-dCTP was removed by extensive buffer exchange with TE buffer (10 mM Tris.HCl pH 8, 0.1 mM EDTA) in a 100 kDa Amicon Ultra spin concentrator (Millipore). The biotinylated DNA was then concentrated to ∼10 mg/ml and stored at −20 °C until needed.

To assemble a DNA-carpet, the DOPC—DOPE-biotin SUV solution was diluted to 1 mg/ml in 500 μl degassed TN150MC buffer (25 mM Tris.HCl pH7.5, 150 mM NaCl, 5 mM MgCl_2_, 0.1 mM CaCl_2_) and warmed to 37 °C. Approximately 300 μl of SUV solution was then infused into a pre-warmed flow cell and incubated at 37 °C for 1 hr. Excess SUVs were washed out with 500 μl warmed, degassed TN150MC buffer at 100 μl/min. 300 μl of a solution of 1 mg/ml neutravidin (Thermo Fisher) in warmed, degassed TN150MC buffer was then infused at a rate of 100 μl/min into the flow cell and incubated at 37 °C for 30 min. Excess neutravidin was washed out with TN150MC buffer as above, and the flow cell infused with 100 μl of a solution containing 1 mg/ml biotinylated sonicated salmon sperm DNA (as prepared above) in warmed, degassed TN150MC buffer and incubated at 37 °C for 30 min. The ports of the flow cell were sealed with parafilm and stored at 4 °C for up to a week.

Prior to use, excess DNA was removed by infusion of 300 μl 0.22 μm filtered and degassed TIRFM buffer (50 mM HEPES.KOH pH 7.5, 300 mM K-glutamate, 50 mM NaCl, 10 mM MgCl_2_, 0.1 mM CaCl_2_, 2 mM DTT, 0.1 mg/ml α-casein, 0.6 mg/ml ascorbic acid, 10% glycerol) with addition of 1 mg/ml α-casein and 1 mM ATPγS and the flow cell incubated at room temperature for 30 min. Conversion of the fluorescence signal detected in TIRF microscopy to the DNA-carpet-bound protein densities was done following the procedure described in the legend of Figure S4 in (Vecchiarelli et al., 2016).

### Fluorescence Recovery After Photobleaching (FRAP)

For FRAP experiments, 488 nm solid-state and 630 nm diode laser was focused to the back focal plane of the objective through an appropriate dichroic mirror (Di01-R405/488/561/635-25×36, Semrock) through the objective lens to illuminate a ∼5 or ∼10 μm (for 488 nm or 630 nm, respectively) diameter spot in the center of the sample area. The laser power was adjusted for ∼80 % bleaching with 5 s exposure for the eGFP or Alexa 647 signals, and four cycles of bleaching/recovery were recorded for each sample and averaged.

### Magnetic Tweezers-based DNA condensation assay

The magnetic tweezer setup and assays conducted with it were performed as previously described (Seol and Neuman, 2011; Seol et al., 2016).

The ability of ParB_F_ to condense *parS_F_*-containing DNA (spDNA) was tested by a custom-built magnetic tweezer setup. In brief, two permanent magnets were used to apply force to micron sized magnetic beads individually tethered to the coverslip of a one inlet flow cell by 5 kb pET28a plasmid-derived DNA tethers. The distance the magnets were held from the beads, and hence the force exerted upon them, was controlled by a linear motor that vertically positions the magnets.

5kb DNA substrates were generated by PCR using either pET28a-*parS_F_* plasmid (for *parS_F_*-containing DNA) or pET28a as templates. pET28-*parS_F_* DNA plasmid was generated by cloning 570 bp DNA segment containing 12 repeats of *parS_F_* native sequence from F-plasmid into pET28a between the BamHI and SphI restriction sites.

Primers used for the PCR contained an extra non-complementary 15 nt at their 5’ ends to encode BsaI restriction sites. The PCR reaction yields a 5.2 kb product incorporating two BsaI restriction sites at its termini. Digestion of this product was followed by ligation with 500 bp DNA “handles” containing either multiple biotin or digitoxin labels. These handles were also generated by Taq-based PCR using pBlueScript II KS as the template, pBlueScript II KS forward (5’-GCT GGG TCT CGG TTG TTC CCT TTA GTG AGG GTT AAT TG) and pBlueScript II KS reverse (5’-GCT GGG TCT CGT GGT TTC CCT TTA GTG AGG GTT AAT TG) primers and either 60 µM biotin-16-dUTP or digoxigenin-11-dUTP (Roche). This results in a 5 kb DNA tether which can be attached to a streptavidin coated magnetic bead at one end and an anti-digoxigenin coverslip surface at the other.

ParB_F_ samples were prepared in modified ATPase buffer (50 mM HEPES.KOH pH 7.5, 100 mM KCl, 5 mM MgCl_2_, 2 mM DTT, 10 mg/ml BSA and 0.1% Tween-20) and infused into a flow cell containing tethered magnetic beads held at 5 pN of force. After the chamber was filled, the flow was stopped, and the force reduced to 0.05 pN. The height of beads was tracked by analysis of diffraction rings generated by illumination of the beads from above and observed with an objective positioned below the flow cell. The extent of condensation by ParB was monitored by the decrease in the height of the beads at 0.05 and 5 pN as compared to controls without protein.

## Acknowledgements

We are grateful to helpful suggestions and discussion of our colleagues Barbara Funnell, David Lane, Michiyo Mizuuchi, Andrea Volante, Min Li, Masaki Osawa, William Carlquist and Shannon Mckie, to Esme Neuman for help in preparation of figures 1 and 7, and to Min Li for help in preparation of Figures 2—figure supplement 3. We thank Stephan Gruber and his colleagues for sharing their findings with us prior to publication. This work was supported by the intramural research fund for National Institute of Diabetes and Digestive and Kidney Diseases (K.M.); the National Heart, Lung, and Blood Institute (K.C.N.), National Institutes of Health, Department of Human Services.

## Figure Supplement Legends

**Source Data Files** (one Excel file with 19 sheets)

Figure 2-Source Data 1:

Numerical data for Figure 2 panels A-E (Microsoft Excel).

Figure 2-Source Data 2:

Numerical data for Figure 2 panels F, G (Microsoft Excel).

Figure 2—figure supplement 2-Source Data 1:

Numerical data for Figure 2—figure supplement 2 (Microsoft Excel).

Figure 2—figure supplement 4-Source Data 1:

Numerical data for Figure 2—figure supplement 4 (Microsoft Excel).

Figure 3-Source Data 1:

Numerical data for Figure 3 panels A-C (Microsoft Excel).

Figure 3-Source Data 2:

Numerical data for Figure 3 panels D (Microsoft Excel).

Figure 4-Source Data 1:

Numerical data for Figure 4 panels A-D (Microsoft Excel).

Figure 4-Source Data 2:

Numerical data for Figure 4 panels E (Microsoft Excel).

Figure 4—figure supplement 1-Source Data 1:

Numerical data for Figure 4—figure supplement 1 (Microsoft Excel).

Figure 4—figure supplement 2-Source Data 1:

Numerical data for Figure 4—figure supplement 2 panels A-D and F (Microsoft Excel).

Figure 5-Source Data 1:

Numerical data for Figure 5 panels A-D (Microsoft Excel).

Figure 5-Source Data 2:

Numerical data for Figure 5 panels E, F (Microsoft Excel).

Figure 5—figure supplement 1-Source Data 1:

Numerical data for Figure 5—figure supplement 1 (Microsoft Excel).

Figure 5—figure supplement 2-Source Data 1:

Numerical data for Figure 5—figure supplement 2 (Microsoft Excel).

Figure 6-Source Data 1:

Numerical data for Figure 6 panel A (Microsoft Excel).

Figure 6-Source Data 2:

Numerical data for Figure 6 panel B-left (Microsoft Excel).

Figure 6-Source Data 3:

Numerical data for Figure 6 panels B-right (Microsoft Excel).

Figure 6-Source Data 4:

Numerical data for Figure 6 panels C (Microsoft Excel).

Figure 6—figure supplement 1-Source Data 1:

Numerical data for Figure 6—figure supplement 1A (Microsoft Excel).

Figure 6—figure supplement 1-Source Data 2:

Numerical data for Figure 6—figure supplement 1B (Microsoft Excel).

Figure 6—figure supplement 2-Source Data 1:

Numerical data for Figure 6—figure supplement 2 (Microsoft Excel).

Figure 6—figure supplement 3-Source Data 1:

Numerical data for Figure 6—figure supplement 3 (Microsoft Excel).

## References

1. Ah-Seng, Y., Lopez, F., Pasta, F., Lane, D., and Bouet, J.Y. (2009). Dual role of DNA in regulating ATP hydrolysis by the SopA partition protein. J Biol Chem 284, 30067–30075.

2. Ah-Seng, Y., Rech, J., Lane, D., and Bouet, J.Y. (2013). Defining the role of ATP hydrolysis in mitotic segregation of bacterial plasmids. PLoS Genet 9, e1003956.

3. Baxter, J.C., and Funnell, B.E. (2014). Plasmid Partition Mechanisms. Microbiol Spectr 2.

4. Bouet, J.Y., Surtees, J.A., and Funnell, B.E. (2000). Stoichiometry of P1 plasmid partition complexes. J Biol Chem 275, 8213–8219.

5. Breier, A.M., and Grossman, A.D. (2007). Whole-genome analysis of the chromosome partitioning and sporulation protein Spo0J (ParB) reveals spreading and origin-distal sites on the Bacillus subtilis chromosome. Mol Microbiol 64, 703–718.

6. Chen, B.-W., Lin, M.-H., Chu, C.-H., Hsu, C.-E., and Sun, Y.-J. (2015). Insights into ParB spreading from the complerx structure of Spo0J and parS. Proc Natl Acad Sci U S A 112, 6613–6618.

7. Davey, M.J., and Funnell, B.E. (1994). The P1 plasmid partition protein ParA. A role for ATP in site-specific DNA binding. J Biol Chem 269, 29908–29913.

8. Davis, M.A., Martin, K.A., and Austin, S.J. (1992). Biochemical activities of the parA partition protein of the P1 plasmid. Mol Microbiol 6, 1141–1147.

9. Ebersbach, G., and Gerdes, K. (2004). Bacterial mitosis: partitioning protein ParA oscillates in spiral-shaped structures and positions plasmids at mid-cell. Mol Microbiol 52, 385–398.

10. Fung, E., Bouet, J.Y., and Funnell, B.E. (2001). Probing the ATP-binding site of P1 ParA: partition and repression have different requirements for ATP binding and hydrolysis. EMBO J 20, 4901–4911.

11. Graham, T.G., Wang, X., Song, D., Etson, C.M., van Oijen, A.M., Rudner, D.Z., and Loparo, J.J. (2014). ParB spreading requires DNA bridging. Genes & development 28, 1228–1238.

12. Hatano, T., Yamaichi, Y., and Niki, H. (2007). Oscillating focus of SopA associated with filamentous structure guides partitioning of F plasmid. Mol Microbiol 64, 1198–1213.

13. Helsberg, M., and Eichenlaub, R. (1986). Twelve 43-base-pair repeats map in a cis-acting region essential for partition of plasmid mini-F. J Bacteriol 165, 1043–1045.

14. Hu, L., Vecchiarelli, A.G., Mizuuchi, K., Neuman, K.C., and Liu, J. (2017). Brownian Ratchet Mechanism for Faithful Segregation of Low-Copy-Number Plasmids. Biophys J 112, 1489–1502.

15. Ivanov, V., and Mizuuchi, K. (2010). Multiple modes of interconverting dynamic pattern formation by bacterial cell division proteins. Proc Natl Acad Sci U S A 107, 8071–8078.

16. Jalal, A.S., Tran, N.T., and Le, T.B. (2020). ParB spreading on DNA requires cytidine triphosphate in vitro. eLife 9, e 53515.

17. Le Gall, A., Cattoni, D.I., Guilhas, B., Mathieu-Demaziere, C., Oudjedi, L., Fiche, J.B., Rech, J., Abrahamsson, S., Murray, H., Bouet, J.Y., and Nollmann, M. (2016). Bacterial partition complexes segregate within the volume of the nucleoid. Nat Commun 7, 12107.

18. Leonard, T.A., Butler, P.J., and Lowe, J. (2005). Bacterial chromosome segregation: structure and DNA binding of the Soj dimer--a conserved biological switch. EMBO J 24, 270–282.

19. Lim, H.C., Surovtsev, I.V., Beltran, B.G., Huang, F., Bewersdorf, J., and Jacobs-Wagner, C. (2014). Evidence for a DNA-relay mechanism in ParABS-mediated chromosome segregation. eLife 3, e02758.

20. Lutkenhaus, J. (2012). The ParA/MinD family puts things in their place. Trends Microbiol 20, 411–418.

21. McLeod, B.N., Allison-Gamble, G.E., Barge, M.T., Tonthat, N.K., Schumacher, M.A., Hayes, F., and Barilla, D. (2017). A three-dimensional ParF meshwork assembles through the nucleoid to mediate plasmid segregation. Nucleic acids research 45, 3158–3171.

22. Modrich, M., and Zabel, D. (1976). Eco RI endonuclease Physical and catalytic properties of the homogeneous enzyme. J Biol Chem 251, 5866–5874.

23. Mori, H., Mori, Y., Ichinose, C., Niki, H., Ogura, T., Kato, A., and Hiraga, S. (1989). Purification and characterization of SopA and SopB proteins essential for F plasmid partitioning. J Biol Chem 264, 15535–15541.

24. Motallebi-Veshareh, M., Rouch, D.A., and Thomas, C.M. (1990). A family of ATPases involved in active partitioning of diverse bacterial plasmids. Mol Microbiol 4, 1455–1463.

25. Murray, H., Ferreira, H., and Errington, J. (2006). The bacterial chromosome segregation protein Spo0J spreads along DNA from parS nucleation sites. Mol Microbiol 61, 1352–1361.

26. Osorio-Valeriano, M., Altegoer, F., Steinchen, W., Urban, S., Liu, Y., and Bange, G. (2019). ParB-type DNA segregation proteins are CTP-dependent molecular switches. Cell 179, 1512–1524.

27. Pillet, F., Sanchez, A., Lane, D., Anton Leberre, V., and Bouet, J.Y. (2011). Centromere binding specificity in assembly of the F plasmid partition complex. Nucleic acids research 39, 7477–7486.

28. Ravin, N.V., Kuprianov, V.V., Gilcrease, E.B., and Casjens, S.R. (2003). Bidirectional replication from an internal ori site of the linear N15 plasmid prophage. Nucleic acids research 31, 6552–6560.

29. Ringgaard, S., van Zon, J., Howard, M., and Gerdes, K. (2009). Movement and equipositioning of plasmids by ParA filament disassembly. Proc Natl Acad Sci U S A 106, 19369–19374.

30. Rodionov, O., Lobocka, M., and Yarmolinsky, M. (1999). Silencing of genes flanking the P1 plasmid centromere. Science 283, 546–549.

31. Sanchez, A., Cattoni, D.I., Walter, J.C., Rech, J., Parmeggiani, A., Nollmann, M., and Bouet, J.Y. (2015). Stochastic Self-Assembly of ParB Proteins Builds the Bacterial DNA Segregation Apparatus. Cell Syst 1, 163–173.

32. Scholefield, G., Whiting, R., Errington, J., and Murray, H. (2011). Spo0J regulates the oligomeric state of Soj to trigger its switch from an activator to an inhibitor of DNA replication initiation. Mol Microbiol 79, 1089–1100.

33. Schumacher, M.A., and Funnell, B.E. (2005). Structures of ParB bound to DNA reveal mechanism of partition complex formation. Nature 438, 516–519.

34. Schumacher, M.A., Piro, K.M., and Xu, W. (2010). Insight into F plasmid DNA segregation revealed by structures of SopB and SopB-DNA complexes. Nucleic acids research 38, 4514–4526.

35. Seol, Y., and Neuman, K.C. (2011). Magnetic tweezers for single-molecule manipulation. Methods Mol Biol 783, 265–293.

36. Seol, Y., Strub, M.P., and Neuman, K.C. (2016). Single molecule measurements of DNA helicase activity with magnetic tweezers and t-test based step-finding analysis. Methods 105, 119–127.

37. Soh, Y.M., Davidson, I.F., Zamuner, S., Basquin, J., Bock, F.P., Taschner, M., Veening, J.W., De Los Rios, P., Peters, J.M., and Gruber, S. (2019). Self-organization of parS centromeres by the ParB CTP hydrolase. Science 366, 1129–1133.

38. Song, D., Rodrigues, K., Graham, T.G.W., and Loparo, J.J. (2017). A network of cis and trans interactions is required for ParB spreading. Nucleic Acids Res 45, 7106–7117.

39. Sugawara, T., and Kaneko, K. (2011). Chemophoresis as a driving force for intracellular organization: Theory and application to plasmid partitioning. Biophysics (Nagoya-shi) 7, 77–88.

40. Taylor, J.A., Pastrana, C.L., Butterer, A., Pernstich, C., Gwynn, E.J., Sobott, F., Moreno-Herrero, F., and Dillingham, M.S. (2015). Specific and non-specific interactions of ParB with DNA: implications for chromosome segregation. Nucleic acids research 43, 719–731.

41. Vecchiarelli, A.G., Han, Y.W., Tan, X., Mizuuchi, M., Ghirlando, R., Biertumpfel, C., Funnell, B.E., and Mizuuchi, K. (2010). ATP control of dynamic P1 ParA-DNA interactions: a key role for the nucleoid in plasmid partition. Mol Microbiol 78, 78–91.

42. Vecchiarelli, A.G., Hwang, L.C., and Mizuuchi, K. (2013). Cell-free study of F plasmid partition provides evidence for cargo transport by a diffusion-ratchet mechanism. Proc Natl Acad Sci U S A 110, E1390–1397.

43. Vecchiarelli, A.G., Li, M., Mizuuchi, M., Hwang, L.C., Seol, Y., Neuman, K.C., and Mizuuchi, K. (2016). Membrane-bound MinDE complex acts as a toggle switch that drives Min oscillation coupled to cytoplasmic depletion of MinD. Proc Natl Acad Sci U S A 113, E1479–1488.

44. Vecchiarelli, A.G., Mizuuchi, K., and Funnell, B.E. (2012). Surfing biological surfaces: exploiting the nucleoid for partition and transport in bacteria. Mol Microbiol 86, 513–523.

45. Vecchiarelli, A.G., Neuman, K.C., and Mizuuchi, K. (2014). A propagating ATPase gradient drives transport of surface-confined cellular cargo. Proc Natl Acad Sci U S A 111, 4880–4885.

46. Watanabe, E., Wachi, M., Yamasaki, M., and Nagai, K. (1992). ATPase activity of SopA, a protein essential for active partitioning of F plasmid. Molecular & general genetics: MGG 234, 346–352.

47. Yamaichi, Y., and Niki, H. (2000). Active segregation by the Bacillus subtilis partitioning system in Escherichia coli. Proc Natl Acad Sci U S A 97, 14656–14661.

